# MUTE Directly Orchestrates Cell State Switch and the Single Symmetric Division to Create Stomata

**DOI:** 10.1101/286443

**Authors:** Soon-Ki Han, Xingyun Qi, Kei Sugihara, Jonathan H. Dang, Takaho A. Endo, Kristen L. Miller, Eun-deok Kim, Takashi Miura, Keiko U. Torii

## Abstract

Precise cell division control is critical for developmental patterning. For the differentiation of a functional stoma, a cellular valve for efficient gas exchange, the single symmetric division of an immediate precursor is absolutely essential. Yet, the mechanism governing the single division event remains unclear. Here we report the complete inventories of gene expression by the Arabidopsis bHLH protein MUTE, a potent inducer of stomatal differentiation. MUTE switches the gene expression program initiated by its sister bHLH, SPEECHLESS. MUTE directly induces a suite of cell-cycle genes, including *CYCD5;1,* and their transcriptional repressors, *FAMA* and *FOUR LIPS*. The architecture of the regulatory network initiated by MUTE represents an Incoherent Type 1 Feed-Forward Loop. Our mathematical modeling and experimental perturbations support a notion that MUTE orchestrates a transcriptional cascade leading to the tightly-restricted, robust pulse of cell-cycle gene expression, thereby ensuring the single cell division to create functional stomata.

**Highlights:**

- Complete inventories of gene expression in stomatal differentiation state are elucidated
- MUTE switches stomatal patterning program initiated by its sister bHLH, SPEECHLESS
- MUTE directly induces cell-cycle genes and their direct transcriptional repressors
- Incoherent feed-forward loop by MUTE ensures the single division of a stomatal precursor

## Introduction

Mirroring the physiological importance of stomatal movement for plant photosynthetic growth, survival, and fitness, a stoma in nearly all land plant species is constituted with a pore surrounded by paired guard cells, which, upon sensing environmental cues, undergo regulated opening and closure (Scarth, 1932). The physiological and evolutionary constraints for forming functional stomata have likely led to a mechanism that strictly enforces a single symmetric cell division to generate stomata with mirror-symmetric, paired guard cells (Chater et al., 2017; Peterson et al., 2010). Yet, the underlying mechanism enabling the only one division event remains unclear.

Stomatal differentiation occurs through stereotypical cell division and differentiation events. In Arabidopsis, an asymmetric entry division of protodermal cell produces a meristemoid, a self-renewing stomatal precursor, which reiterates asymmetric amplifying divisions but eventually differentiate into a guard mother cell (GMC). Stomatal differentiation completes following the single symmetric cell division (SCD) of the GMC (Han and Torii, 2016; Lau and Bergmann, 2012). Three basic-helix-loop-helix (bHLH) proteins, SPEECHLESS (SPCH), MUTE, and FAMA drive the sequential steps of stomatal differentiation (Han and Torii, 2016; Lau and Bergmann, 2012). They are expressed transiently in a respective developmental window, and their loss-of-function mutations result in a failure to initiate stomatal cell-lineages (*spch*), arrested meristemoids (*mute*), and multinumeral GMC-like tumors (*fama*), respectively (MacAlister et al., 2007; Ohashi-Ito and Bergmann, 2006; Pillitteri et al., 2007). SPCH, MUTE, and FAMA form a heterodimer with bHLH proteins, SCREAM (SCRM, also known as ICE1) and SCRM2 (Kanaoka et al., 2008). In addition, Myb genes *FOUR LIPS* (*FLP*) and *Myb88* restrict GMC-divisions, and *flp myb88* mutants exhibit multinumeral GCs (Lai et al., 2005).

Extrinsic cell-cell signaling enforces proper stomatal patterning through inhibition of these stomatal bHLH proteins. A small secreted peptide EPIDERMAL PATTERNING FACTOR2 (EPF2) is perceived by the receptor kinase ERECTA and its partner TOO MANY MOUTHS (TMM), and the signal inhibits SPCH (Hara et al., 2009; Hunt and Gray, 2009; Lee et al., 2012; Nadeau and Sack, 2002). SPCH and SCRM directly induce *EPF2* and *TMM* expression, thereby constituting a negative feedback loop (Horst et al., 2015; Lau et al., 2014). During the meristemoid-to-GMC transition, EPF1 perceived by ERECTA-LIKE1 (ERL1) enforces stomatal spacing (Hara et al., 2007; Lee et al., 2012; Qi et al., 2017). Here, MUTE directly induces *ERL1*, which mediates an autocrine signaling for proper GMC differentiation (Qi et al., 2017)

Plants possess a large number of cyclin genes, likely reflecting their continuous development throughout the life cycle (De Veylder et al., 2007; Scofield et al., 2014). Among them, D-type cyclins (CYCDs) associate with CYCLIN DEPENDENT KINASE A1;1 (CDKA1;1) to drive G1/S-phase transition, whereas A-type cyclins (CYCAs) complex with CDKA for S/G2 transition as well as with CDKB1;1 to suppress endoreduplication (Boudolf et al., 2009; De Veylder et al., 2007; Scofield et al., 2014). Specific roles of core-cell cycle regulators have been associated with stomatal development. For example, CYCD4 is involved in stomatal-lineage divisions in the hypocotyls (Kono et al., 2007). Dominant-negative form of CDKB1;1 and CDKA1;1, as well as higher-order mutants of CYCAs inhibit the SCD of GMCs (Boudolf et al., 2004; Yang et al., 2014). It was reported that FLP directly represses *CDKB1;1* and *CDKA1;1* (Xie et al., 2010; Yang et al., 2014), and FAMA binds to the promoter region of *CDKB1;1* (Hachez et al., 2011). Although these studies demonstrated the repressive roles for FAMA and FLP in the GMC division, it is not clear how the single symmetric division event is initiated and robustly controlled.

Among the stomatal transcription factors, *MUTE* is the only one whose overexpression confers constitutive differentiation of stomata with paired GCs (Pillitteri et al., 2008; Pillitteri et al., 2007; Trivino et al., 2013). Overexpression of *SPCH* and *FAMA* confers highly-divided stomatal-lineage cells and singular GCs, respectively (MacAlister et al., 2007; Ohashi-Ito and Bergmann, 2006). We thus hypothesized that MUTE governs the gene regulatory networks to create stomata. To test this, we performed a genome-wide profiling of early MUTE-responsive genes.

Comparison of SPCH and MUTE target genes revealed how MUTE disconnects stomatal-lineage cells from extrinsic inhibitory signals, thus ‘locks in’ the differentiation program. Contrary to the known role of MUTE in terminating the proliferating meristemoids (Pillitteri et al., 2007), our study identified cell cycle and cell division genes as overwhelming majorities of the MUTE targets. MUTE directly binds to the promoters and upregulates novel and previously described cell-cycle regulators of the GMC symmetric division process. At the same time, MUTE directly binds to the promoters and upregulates *FAMA* and *FLP*, which in turn repress the cell-cycle regulators. Our mathematical modeling predictions and experimental perturbations of network motif demonstrate that an incoherent feed-forward loop mediated by MUTE, cell cycle regulators, and FAMA/FLP is sufficient to articulate the single symmetric division event with high fidelity. Our study establishes the role for MUTE in orchestrating a transcriptional cascade leading to stomatal differentiation and defines a core regulatory circuit for the single symmetric division.

## Results

### MUTE induces and represses specific sets of transcriptomes

To obtain the most complete inventories of transcriptional changes driven by MUTE, we employed an estradiol-inducible *MUTE* overexpressor (*iMUTE*) line (Qi et al., 2017), which upon estradiol treatment, triggered a rapid, 200-fold increase in *iMUTE* transcripts within two hours, reaching to >1600 fold increase in 12 hours (Figure S1A). The seedlings eventually differentiated epidermis solely composed of stomata, many expressing the mature GC marker E994 (Figure 1A-D). GC differentiation is governed by the sister gene *FAMA* (Hachez et al., 2011). In order to identify the direct MUTE targets and not those governed by FAMA, we examined the transcriptional changes before *FAMA* expression peaks. We performed paired-end sequencing of RNA from *iMUTE* and un-induced controls (see Methods and Table S1). The three biological replicates showed high reproducibility, with the Pearson’s correlations between log RPKM, >0.980 among samples of the same condition (Figure S1B).

**Figure 1.**
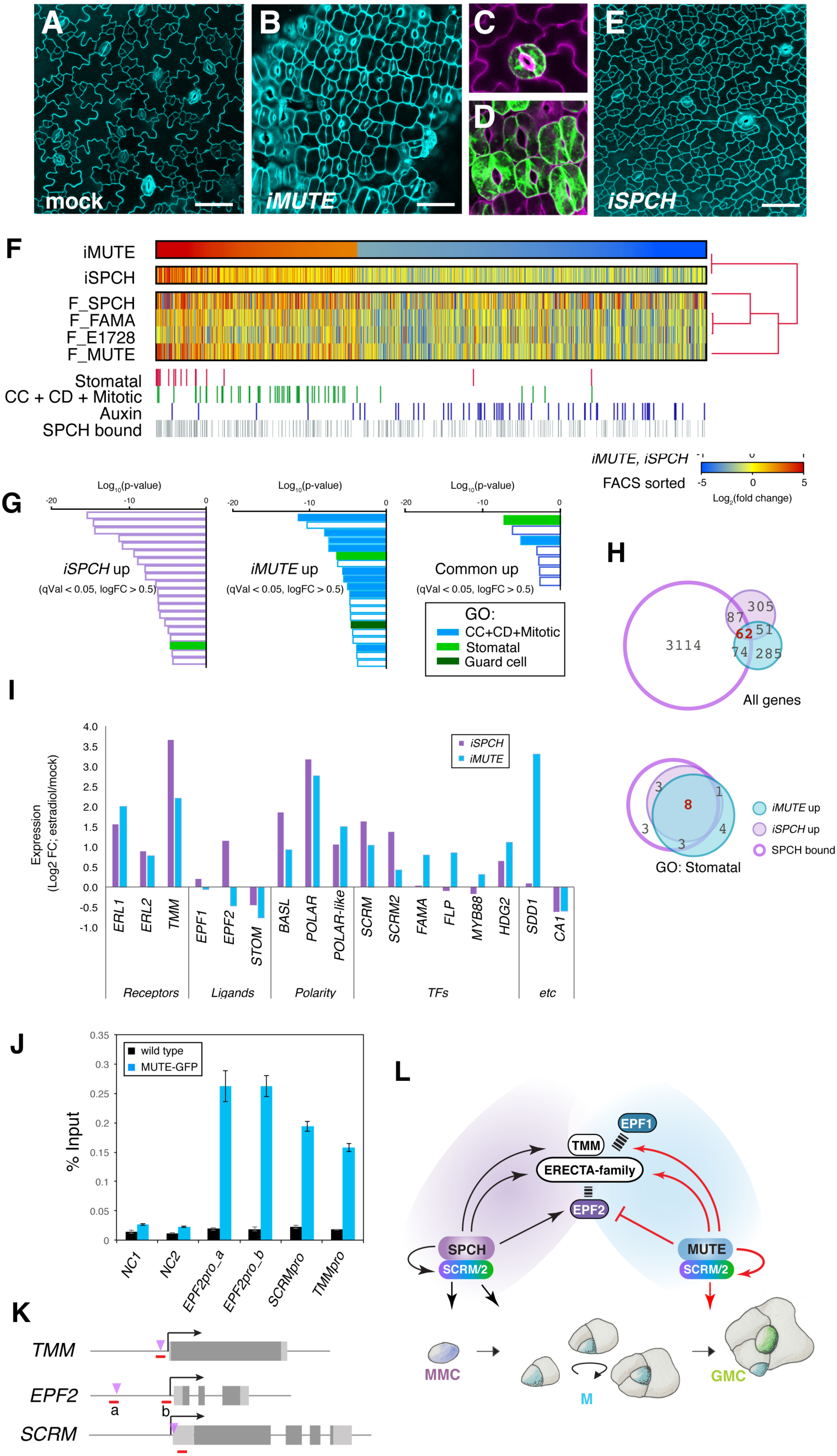
Transcriptomic profiling of MUTE target genes reveals a framework of stomatal cell-state switch. (A-E) Epidermal phenotypes of 3-day-old seedlings carrying inducible *MUTE* construct, either mock treated (A, C) or estradiol-induced (*iMUTE*) in media (B, D). Mature stomata of mock (C) and *iMUTE* (D) cotyledon epidermis expressing GC GFP marker E994. Induced overexpression of *SPCH* (*iSPCH*) showing excessive epidermal cell divisions (E). Scale bars, 50 μm. (F) Heat map of *iMUTE* DEGs (log_2_ FC ≥ 0.5 and ≤ -0.5, respectively, q-val ≤ 0.05) by RNA-seq analysis. Their expression fold-changes by *iSPCH* (Lau et al., 2014) as well as in FACS-sorted stomatal-lineage cells (Adrian et al., 2015) are shown as heat maps below. Genes in GO categories: red, “stomatal”; green, “cell cycle, cell division and mitotic (CC+CD+Mitotic)”; blue, “auxin”; and gray, SPCH-bound according to published ChIP-seq data (Lau et al., 2014). (G) GO categories of top *iSPCH* up, *iMUTE* up, and common up (log_2_ fold change ≥ 0.5, q-val ≤ 0.05), ranked by p-values. Green, “stomatal”; blue, “CC+CD+Mitotic”, and dark green, “guard cells”. For complete lists, see Figure S2 and Table S2. (H) Venn diagrams of *iMUTE*-up (light sky blue), *iSPCH-*up (lilac), and SPCH-bound genes (purple outline) for all genes (top) and for the combined GO categories “stomatal” (bottom). (I) Expression FC of known stomatal regulators by *iMUTE* (blue) and *iSPCH* (purple). (J) ChIP assays showing specific binding of functional MUTE-GFP at the promoter regions. NC, Negative Control; NC1, 5’ intergenic region of *ACTIN2*; NC2, promoter region of *AGAMOUS*. Bars, average of three technical repeats. Error bars, s.e.m. See additional two biological replicates in Figure S4. (K) Gene structures. Light gray rectangles, UTRs; dark gray rectangles, exons; arrows, transcriptional start sites; red line, amplicons; purple triangle, known SPCH binding sites (Lau et al., 2014). (L) Updated model of stomatal cell-state switch by MUTE. See main text for detail.

472 *iMUTE* upregulated genes and 818 downregulated genes were extracted after q value =0.05 cutoff and Log_2_ FC >0.5 (Figure 1F, Table S1). The Gene Ontology (GO) categories for the *iMUTE* upregulated genes are overwhelmingly specific to cell cycle, cell division, and mitosis (CC+CD+Mitotis), including cell cycle (GO:00070429), cell division (GO:0051301), DNA replication (GO:0006260), and mitotic nuclear division (GO:0000280). Highly enriched categories also include stomatal complex morphogenesis (GO:0010103), guard cell differentiation (GO:0010052) and microtubule-based movement (GO:0007018)(Figures 1G, S2, Table S2). In contrast, the GO enriched categories for *iMUTE* downregulated genes include response to auxin (GO:0009733), auxin polar transport (GO:0009926), and cell-wall loosening (GO:0009828)(Figures 1F, S2, Table S2). The clear categorizations of specific *iMUTE-*regulated genes indicate that MUTE promotes cell cycle/cell division and stomatal development, while repressing non-stomatal cell characters. A comparison of *iMUTE* differentially expressed genes (DEGs) with published fluorescent assisted cell-sorted (FACS) stomatal-lineage transcriptomes (Adrian et al., 2015) suggests that *iMUTE* DEGs share a similarity with those enriched in *MUTE-* expressing cells (Figures 1F, S1E). Low reproducibility among the FACS-sorted samples, however, limited the reliable comparisons (Figure S1D; see Methods).

### MUTE acts as a transcriptional switch for the stomatal patterning ligand-receptor systems

The closest paralog of *MUTE, SPCH* drives the initial entry into stomatal cell lineages, and its induced overexpression (*iSPCH*) results in epidermis with excessive asymmetric divisions (MacAlister et al., 2007; Pillitteri et al., 2007)(Figure 1E). To understand the extent of their shared and specific functions, we next compared their target genes. An induced SPCH (*iSPCH*) RNA-seq analysis was performed essentially at the same condition (8 hours of induction using 4-day-old seedlings) (Lau et al., 2014), thus we re-analyzed the published *iSPCH* data for direct comparison. 24% (113/472) of *iMUTE* upregulated genes are shared by *iSPCH* (Figure 1F, H, Table S1). To further correlate their shared transcriptional response to physical genome-wide SPCH binding locations (Lau et al., 2014), we analyzed the extent by which the promoters of the co-regulated genes are occupied by SPCH. 55% (62/113) of SPCH and MUTE co-upregulated genes are bound by SPCH, whereas only 22% (87/392) of SPCH-specific upregulated genes are bound by SPCH (Figure 1F, H, Table S1). Thus, extracting the transcriptional response shared by SPCH and MUTE highly enriches the selected set of the SPCH direct targets.

The most highly enriched GO-category for *iSPCH* and *iMUTE* shared co-upregulated genes is stomatal complex development, where 74% (14/19) of genes are SPCH-bound (Figures 1G, S2, Table S3). All eight *iMUTE*-up/*iSPCH*-up/SPCH-bound genes are known players of stomatal development: *TMM, ERL1, ERL2, BASL*, *POLAR,* and *POLAR-LIKE*, *SCREAM* (*SCRM*), and *HDG2* (Figure 1G, H, I, Table S3). *SCRM2* was also co-upregulated (Log_2_=0.43, qVal=3.94E-14)(Figure 1I, Table S1). A subsequent qRT-PCR analysis confirmed their induction (Figure S3). On the other hand, *CARBONIC ANHYDRASE1* (*CA1*), which mediates inhibition of stomatal development at elevated CO_2_ levels (Engineer et al., 2014), and *STOMAGEN* were repressed by both *iSPCH* and *iMUTE* (Figure 1I, Table S1).

To rule out the possibility that *iMUTE* causes a non-specific, promiscuous induction of SPCH targets, we further tested whether the promoters of these genes are indeed occupied by the functional MUTE protein expressed transiently during the meristemoids-to-GMC transition. For this purpose, chromatin-Immunoprecipitation (ChIP) assays were performed using Arabidopsis *mute* plants complemented by the functional MUTE-GFP protein driven by its own promoter (*proMUTE::MUTE-GFP*) using the *scrm-D* enabled background (Horst et al., 2015; Pillitteri et al., 2007; Qi et al., 2017). Indeed, direct associations of MUTE with the promoters of *TMM, SCRM,* as well as *BASL* and *POLAR* were detected, indicating that they are the *bona fide*, direct MUTE targets (Figures 1J, S4). The strong MUTE-GFP association was detected within the location of known SPCH-binding sites, many possessing an E-box, which is a known bHLH binding sites (Figures 1K, S4, Table S3).

*EPF2* is a known direct target of SPCH (Horst et al., 2015; Lau et al., 2014). Although *iSPCH*, triggered >30 fold increase in *EPF2* expression, *iMUTE* directly repressed *EPF2* (-0.47, qVal=1.50E-02)(Fig. 1I, Fig. S3). The ChIP assays detected the direct MUTE binding to the SPCH binding site within the *EPF2* promoter (Figures 1J, K, S4), indicating that MUTE changes transcription of EPF2 via replacing SPCH. On the other hand, both *iSPCH* and *iMUTE* directly upregulate the receptors, *ERLs* and *TMM* (Figures 1I, J, K, S3, S4)(Horst et al., 2015; Lau et al., 2014; Qi et al., 2017). It is worth noting that EPF1, the sister peptide of EPF2, is perceived by ERL1 and TMM to regulate MUTE activity during the GMC differentiation (Qi et al., 2017). Together, our findings suggest that MUTE acts as a transcriptional switch for the stomatal patterning ligand-receptor system, eliminating the earlier signal EPF2 induced by SPCH, while maintaining the expression of shared receptors to perceive the later signal, EPF1 (Figure 1L).

### MUTE directly upregulates cell-cycle genes driving the symmetric division of stomata

*MUTE* terminates asymmetric division of meristemoids (Pillitteri et al., 2007). Unexpectedly, the most significantly *iMUTE*-upregulated genes belong to the combined GO-categories CC+CD+Mitotic, suggesting that MUTE is a potent inducer of cell division (Figures 1, F, G, S2, Table S2). Indeed, 46 genes belonging to the CC+CD+Mitotic categories are up by *iMUTE*, whereas only 2 genes are specifically up by *iSPCH*, which initiates stomatal-lineage divisions (Figures 1A, 2B, Table S3). Among the cell cycle genes (Figures 2A, S5), *iMUTE* strongly induced B-type Cyclin-Dependent Kinase genes *CDKB1;1*, and *CDKB2;1* as well as A-type cyclins, most notably *CYCA1;1, CYCA2;2*, and *CYCA2;3* (Figures 2B, S5). A subsequent ChIP analysis demonstrated that functional MUTE protein associates with the promoter regions of these cell-cycle regulator genes, indicating that they are direct MUTE targets (Figures 2C, D, S4). qRT-PCR time-course analyses confirmed the increase of *CDKB1;1, CYCA2;2,* and *CYCA2;3* transcript levels ∼4 hrs after *iMUTE* induction (Figure S5). CDKB1;1 and CYCA2s are known to promote the GMC symmetric division (Boudolf et al., 2004; Xie et al., 2010) and, consistently, they are enriched in *MUTE*-expressing transcriptome (Adrian et al., 2015)(Figure 2B). Our results suggest that MUTE promotes the SCD of GMCs through direct upregulation of *CDKB1;1* and *CYCA2s*.

**Figure 2.**
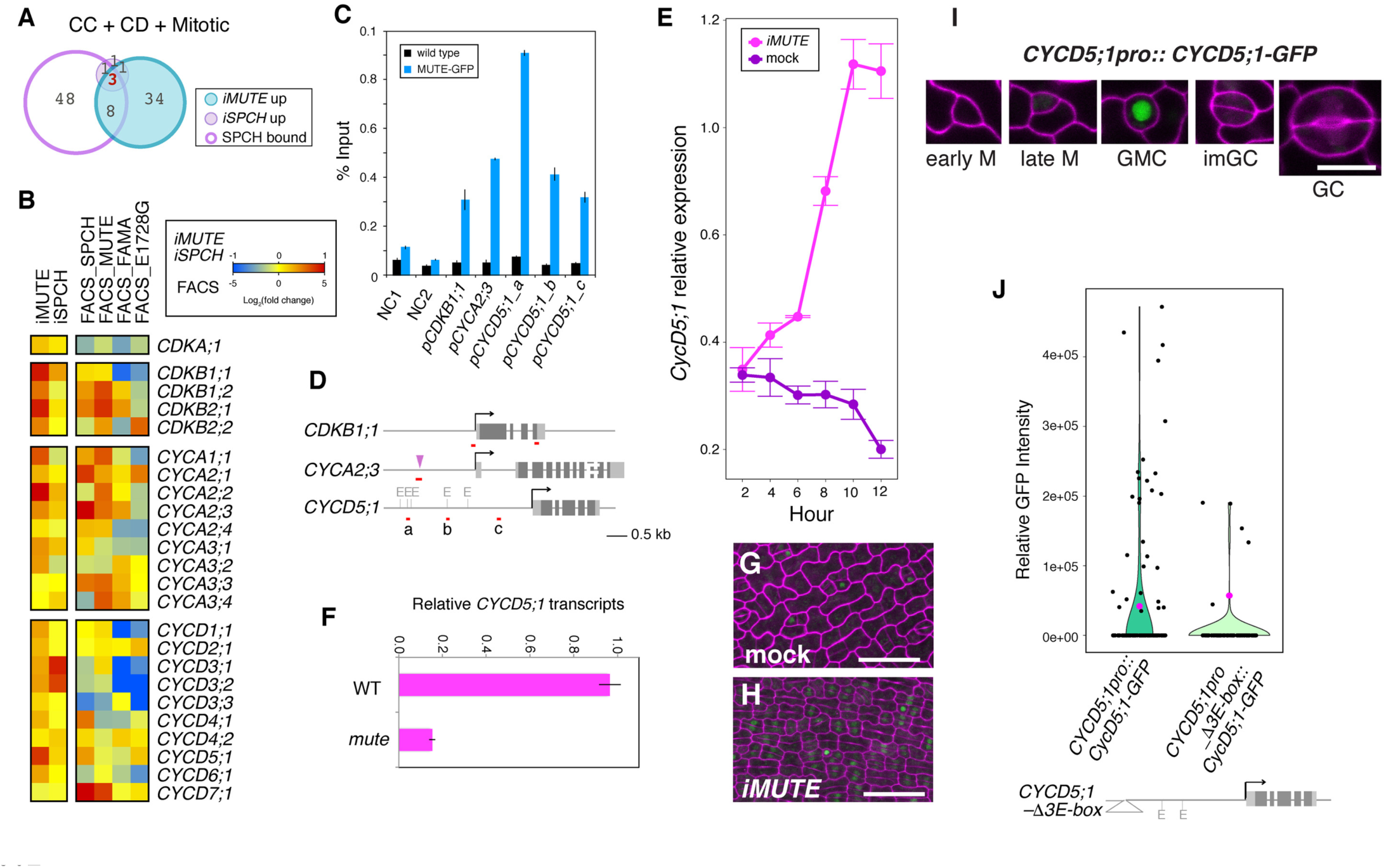
Direct role of MUTE in promoting cell-cycle gene expression. (A) Venn diagrams of *iMUTE*-up (light sky blue), *iSPCH-*up (lilac), and SPCH-bound genes (purple outline; left) for the combined GO categories CC+CD+Mitotic(right). For gene lists, see Table S2. (B) Heat map showing expression (Log_2_ FC) of cell cycle genes by *iMUTE*, *iSPCH*, and FACS-sorted stomatal-lineage cells. (C) ChIP assays showing the binding of functional MUTE-GFP at the promoter regions. Bars, average of three technical repeats. Error bars, s.e.m. See additional two biological replicates in Figure S4. (D) Diagrams of *CDKB1;1*, *CYCA2;3* and *CYCD5;1* loci. Light gray rectangles, UTRs; dark gray rectangles, exons; arrows, transcriptional start sites; red line, amplicons; purple triangle, known SPCH binding sites (Lau et al., 2014); E, E-boxes. (E) *iMUTE* triggers rapid induction of *CYCD5;1* transcripts. Shown are time course of CYCD5;1 relative expression by *iMUTE* vs. mock, normalized against *ACTIN* (*ACT2*). Dots, mean values of three technical replicates; error bars, s.d. For all three biological replicates, see Figure S5. (F) Relative *CYCD5;1* expression in 8-day-old wild-type (WT) and *mute* seedlings normalized against *ACT2*. Bars, mean of three technical replicates. Error bars, s.e.m. For additional two biological replicates, see Figure S5. (G and H) *iMUTE* triggers ectopic overexpression of *CYCD5;1-GFP* on developing epidermis. Mock (G) and *iMUTE* (H) for 40 hours of germination. Scale bars, 50 μm. (I) *CYCD5;1pro::CYCD5;1-GFP* in stomatal lineage cells. Images are taken under the same magnification. Scale bar, 10 μm. (J) Violin plots of relative GFP intensity within nuclei of meristemoids from 12-day-old seedlings expressing *CYCD5;1pro::CYCD5;1-GFP* (left; n=122) and *CYCD5;1pro_⊗3E-box::CYCD5;1-GFP* (right; n=86), whereby the three E-boxes in the amplicon a (D) are removed (bottom). Black dots, values from individual nuclei; pink dots, means. p-value, Wilcoxon Rank Sum Test.

### CYCD5;1, a direct MUTE target, accumulates transiently prior to the GMC symmetric division

The previous stomatal-lineage transcriptome study reported *CYCD7;1* as a GMC-specific D-type cyclin (Adrian et al., 2015). However, our RNA-seq and time-course transcript analyses showed that *MUTE* induction has no effects on the expression of *CYCD7;1* (Log_2_FC =0.07, qVal=1.00E+00; Figures 2B, S5, Table S1). Therefore, MUTE does not activate *CYCD7;1* expression. In contrast, *iMUTE* strongly induces *CYCD5;1,* which has not been associated with stomatal development previously (Figure 2B). The functional MUTE protein robustly binds to the promoter of *CYCD5;1* (Figure 2C, D), and *iMUTE* rapidly induces *CYCD5;1* transcripts (Figure 2E), demonstrating that *CYCD5;1* is a MUTE target. Consistently, *CYCD5;1* transcript levels were substantially reduced in the *mute* mutant background (Figures 2F, S4).

To understand the role of *CYCD5;1* in stomatal development, we generated Arabidopsis plants expressing CYCD5;1-GFP driven by its own promoter (*CYCD5;1pro::CYCD5;1-GFP*). A strong signal was detected within the nuclei of a GMC (Figure 2G, I). Consistently, *iMUTE* vastly increased the cells accumulating CYCD5;1-GFP (Figure 2H). The *CYCD5;1* promoter possesses five E-boxes, 3 of which situate where the robust in vivo MUTE-binding is detected (Figure 1C, D; Amplicon **a**). Deletion of the 3 E-boxes diminished though not abolished the GFP signals in meristemoids (Figure 1J), supporting that MUTE upregulates *CYCD5;1* via direct binding to its promoter.

To address the accumulation dynamics of CYCD5;1 during stomatal differentiation, we next performed time-lapse live imaging using the double transgenic lines expressing CYCD5;1-GFP and plasma-membrane RFP (Figure 3A; Movie S1). CYCD5;1-GFP accumulates in the nucleus of a meristemoid within 3-4 hrs (3.3 ± 1.4 hrs, n=25) after the last asymmetric division, reaches maximum ∼10 hrs (10.6 ± 4.1 hrs, n=26), and disappears ∼8 hrs (7.9 ± 1.1 hrs, n=28) in prior to the symmetric division (Figure 3A, Movie S1). Consistent with our finding that MUTE directly activates *CYCD5;1* expression, functional MUTE-GFP accumulates in the nucleus of a meristemoid ∼1.5 hrs (1.6 ± 0.4 hrs, n=25) after the last asymmetric division, thus preceding the accumulation of CYCD5;1-GFP (Figure 3B, Movie S2). Functional FAMA-GFP, which restricts the SCD, appears ∼3.5 hrs (3.3 ± 0.4 hrs n=25) prior to the symmetric division (Fig 3C, Movie S3). Together, our time-lapse studies elucidate the in vivo dynamics of CYCD5;1 within the narrow developmental window between MUTE and FAMA expression, suggesting its role in the GMC symmetric division.

**Figure 3.**
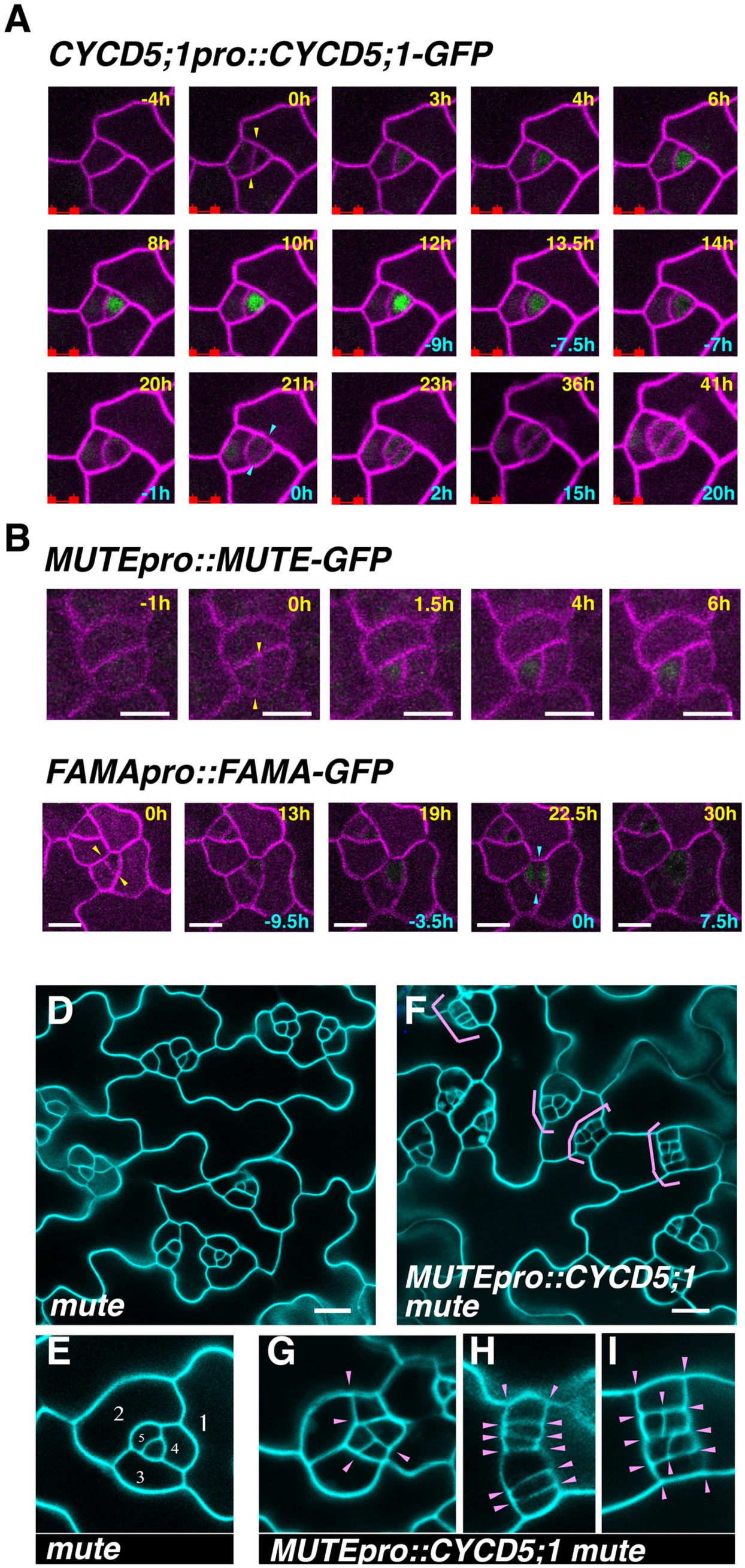
CYCD5;1 promotes GMC symmetric cell division. (A) Transient CYCD5 accumulation (*CYCD5;1pro::CYCD5;1-GFP)* during meristemoid-to-GMC transition revealed by the time-lapse imaging. Yellow arrowheads indicate the last amplifying asymmetric cell division (ACD) of a meristemoid (hour 0 in yellow). Cyan arrowhead indicates the single symmetric cell division (SCD) that gives rise to paired guard cells (hour 0 in cyan). Scale bar, 10 μm. For a full sequence, see Movie S1. (B) MUTE (*MUTEpro::MUTE-GFP*) accumulation dynamics revealed by the time-lapse imaging. MUTE-GFP accumulates immediately after the last ACD (yellow arrowheads, hour 0), preceding the accumulation of CYCD5;1 (A). Scale bar, 5μm. For a full sequence, see Movie S2. (C) FAMA (*FAMApro::FAMA-GFP*) accumulation dynamics revealed by the time-lapse imaging. FAMA-GFP accumulations are visible ∼4 hours before the SCD of GMC (cyan arrowheads, hour 0). Scale bar, 10 μm. For a full sequence, see Movie S3. (D-I) *CYCD5;1* expression in the arrested *mute* meristemoids is sufficient to trigger SCD-like divisions. Shown are cotyledon epidermis images from 2-week-old *mute* (D; inset E) and *mute* expressing *proMUTE::CYCD5;1* (F; insets G-I). In *mut*e, each meristemoid arrests after ACDs in an inward-spiral manner (A and E, numbered by the order). *MUTEpro::CYCD5;1* in *mute* confers aberrant divisions (F, pink brackets) in perpendicular or parallel orientations (pink arrowheads, G-I). Scale bars, 20 μm.

### *CYCD5;1* is sufficient to trigger symmetric division-like divisions of arrested *mute* meristemoids

To address whether *CYCD5;1* expressed in the meristemoids is sufficient for the symmetric division in the absence of *MUTE*, we expressed *CYCD5;1* driven by the *MUTE* promoter into *mute* null mutant background (Figure 3D-I). The *mute* meristemoids undergo several rounds of spiral asymmetric divisions and arrest (Pillitteri et al., 2007)(Figure 3D, E). Strikingly, *MUTEpro::CYCD5;1* triggered occasional aberrant divisions of arrested *mute* meristemoid (Figure 3F-I, pink brackets). These aberrant divisions are in parallel or perpendicular orientations (Figure 3G-I, pink arrowheads), resembling to GMC-tumors seen in *fama* (Ohashi-Ito and Bergmann, 2006). Therefore, the expression of *MUTEpro::CYCD5;1* alone could drive symmetric-cell-division-like divisions of arrested *mute* meristemoids. None of these aberrantly-divided tumors differentiated into stomata, consistent with the notion that other MUTE target genes are necessary for guard cell differentiation programs.

### *FAMA* and *FLP*, which restrict the symmetric division, are also direct MUTE targets

FAMA and FLP restrict the SCD of a GMC through direct binding to the promoters of *CDKB1;1* (Hachez et al., 2011; Xie et al., 2010). In addition, FLP directly suppresses *CDKA1;1* expression (Xie et al., 2010; Yang et al., 2014). It remains unknown, however, what triggers their expressions. To address this question, we investigated the regulatory relationships of MUTE with FLP and FAMA. Our RNA-seq analysis identified both *FAMA* and *FLP* as *iMUTE-*specific upregulated genes, not upregulated by *iSPCH* (Figure 1I, Table S1). Both *FAMA* and *FLP* expressions are induced at around 8 hours after the *iMUTE* induction (Figures 4A; S3), slightly delayed from early stomatal genes and cell-cycle genes (Figures S3, S5). The *FAMA-GFP* signal was not detected in the *mute* epidermis (Figure 4B). On the other hand, the previously-reported *FLP* reporter, *FLPpro::GUS-GFP* (Lai et al., 2005), exhibited a basal expression throughout leaf epidermis with stronger signals in GMCs (Figure 4B). The basal *FLP* reporter signal persisted in arrested *mute* meristemoids (Figure 4B). Consistently, in *mute*, FAMA transcripts are at a detection limit whereas FLP transcripts were substantially reduced yet detectable (Figures 4C, S4). Thus, MUTE is absolutely required for *FAMA* expression, while MUTE boosts *FLP* expression during the GMC transition.

**Figure 4.**
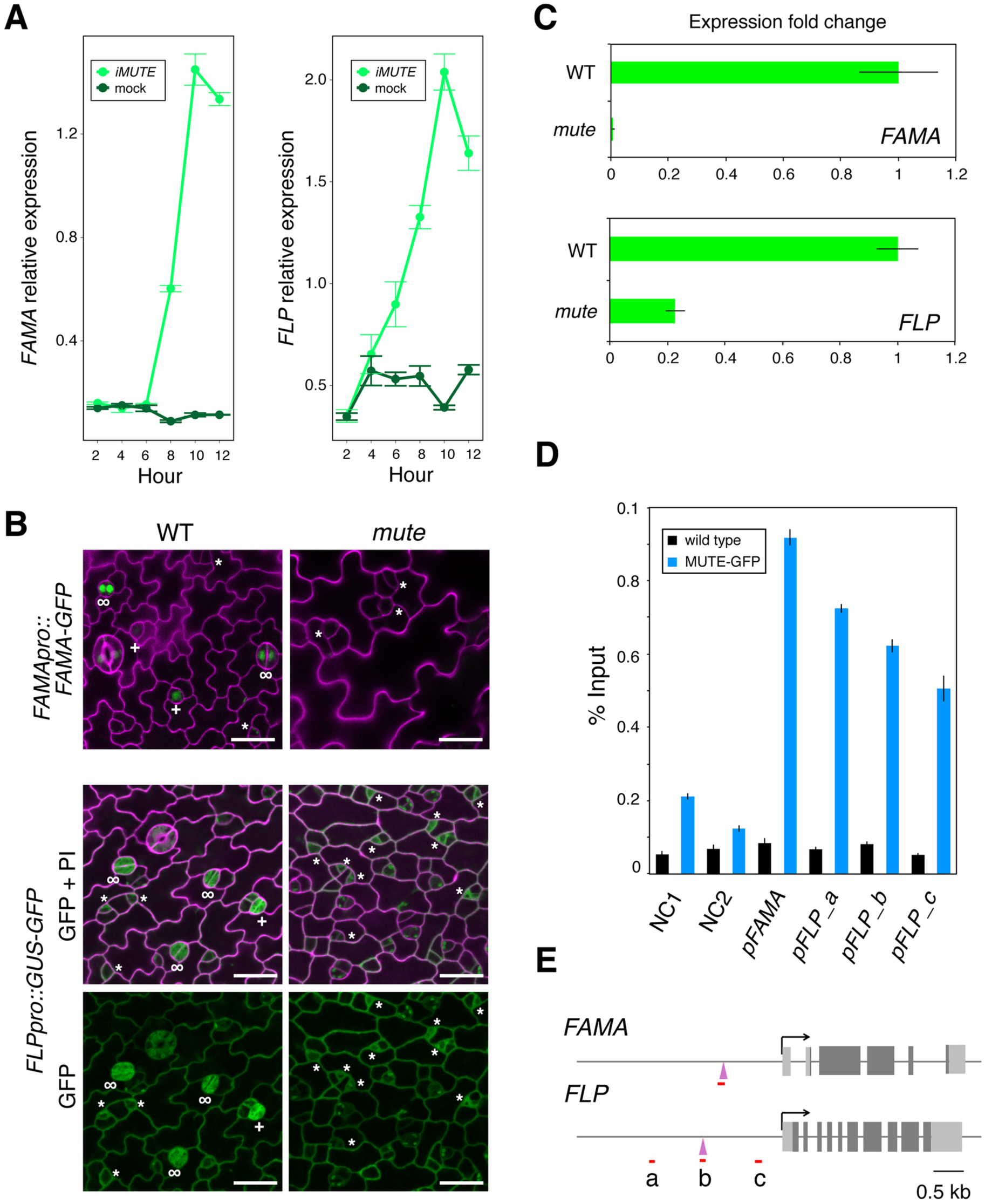
*FAMA* and *FLP* are direct targets of MUTE. (A) Time course of *FAMA* and *FLP* relative expression by *iMUTE* vs. mock, qRT-PCR normalized against *ACT2*. Dots, mean values of three technical replicates; error bars, sem. For all three biological replicates, see Figure S3. (B) *FAMApro::FAMA-GFP* and *FLPpro::GUS-GFP* reporter expression in 5-day-old wild-type and *mute* epidermis. Asterisks, meristemoids; plus, GMCs; infinity, immature GCs. Scale bars, 20 μm. (C) *FAMA* and *FLP* expression fold change in 7-day-old wild-type (WT) and two biological replicates of *mute* seedlings normalized against *ACTIN*. Bars, mean of three technical replicates. Error bars, s.d. See additional four biological replicates in Figure S5. (D) ChIP assays showing the binding of functional MUTE-GFP at *FAMA* and *FLP* promoter regions. For detail see Figure 1. See additional two biological replicates in Figure S5. (E) Diagrams of *FAMA* and *FLP* loci. Light gray rectangles, UTRs; dark gray rectangles, exons; arrows, transcriptional start sites; red line, amplicons; purple triangle, known SPCH binding sites (Lau et al., 2014).

We subsequently tested whether *FAMA* and *FLP* are direct MUTE targets. ChIP experiments with MUTE-GFP detected strong signals at the *FAMA* and *FLP* promoters, overlapping with the regions of known SPCH binding peaks (Figures 4D,E, S4). Note, however, that *iSPCH* does not upregulate their expressions (Figure 1F, Table S1), suggesting that MUTE takes over the binding sites from SPCH and, in this case, activates the expression of *FAMA* and *FLP* to achieve stomatal differentiation.

### Incoherent type 1 feed forward loop (I1-IFF) ensuring the single symmetric cell division

Our study revealed that MUTE directly upregulates both cell cycle regulators that positively drive SCD and transcription factors that negatively regulate SCD via direct repression of the cell-cycle genes. In the field of network dynamics and behaviors, the MUTE-driven transcriptional network motif represents a typical Incoherent Type I Feed-Forward Loop (I1-FFL) constituted by the three nodes, X, Y, and Z (Mangan and Alon, 2003). Here, MUTE (X) situates at the top of the network, which upregulates both *FAMA*/*FLP* (Y) as well as *CDKs/CYCs* (Z), whereas FAMA and FLP (Y) directly repress *CDKs/CYCs* (Z)(Figure 5A). The I1-FFL is known to produce a single highly-tuned pulse of output Z (Mangan and Alon, 2003), in this case the cell-cycle regulators. To elucidate if the I1-FFL orchestrated by MUTE is necessary and sufficient for the transient expressions and activities of the network output Z (CDKs/CYCs) during stomatal development, we took both mathematical modeling and experimental approaches.

**Figure 5.**
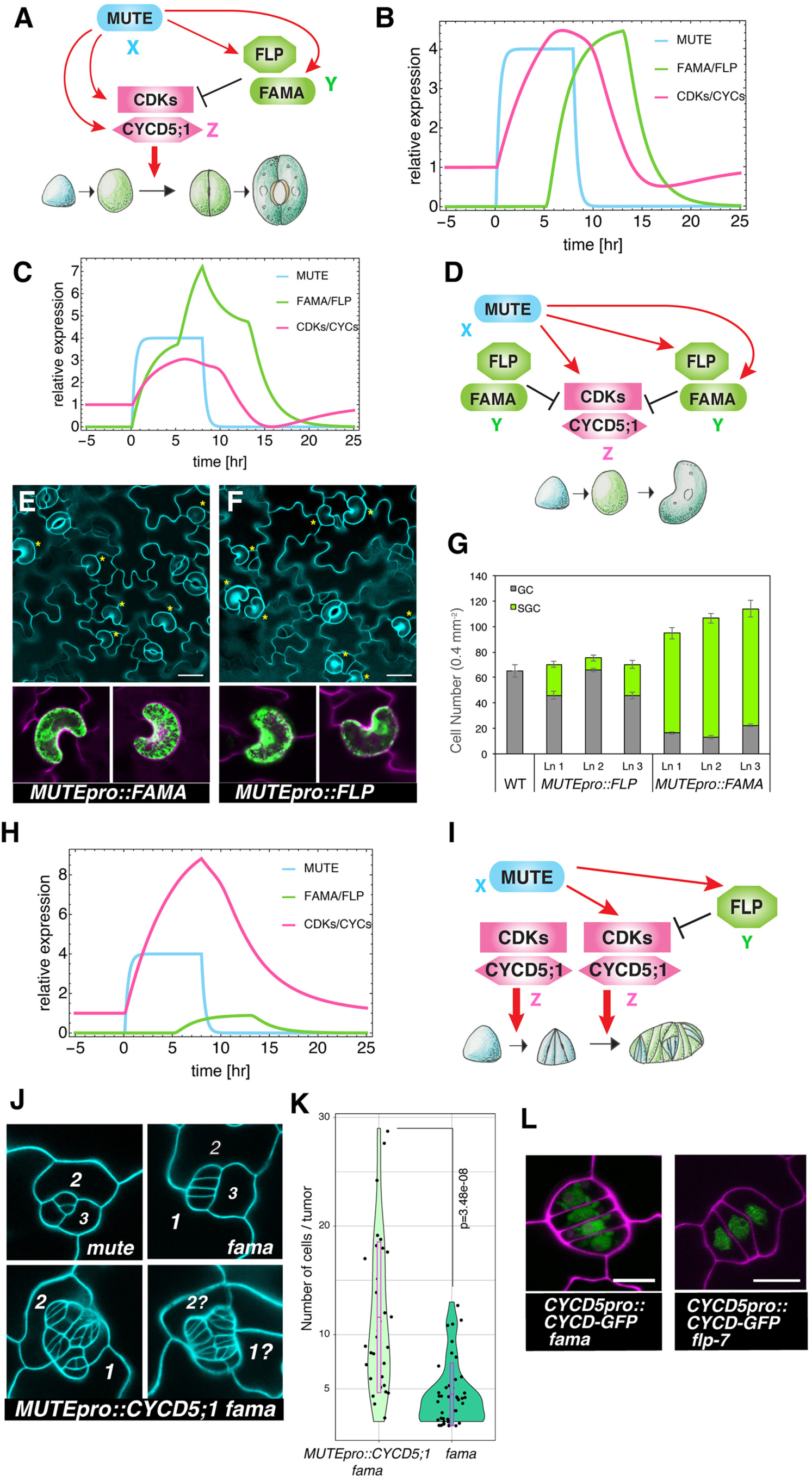
MUTE orchestrates a single symmetric cell division to produce a stoma with paired guard cells. (A) Architecture of type I incoherent feed-forward loop (I1-FFL) orchestrated by MUTE. MUTE induces both cell-cycle regulators driving SCD and transcription factors that repress these cell-cycle regulators. (B) Dynamics of MUTE, FAMA/FLP and CDKs/CYCs reproduced by the mathematical model. Pulse of CDKs/CYCs is generated by successive induction by MUTE and repression by FAMA/FLP. (C) Mathematical modeling prediction of the effects of precocious expression of FAMA/FLP. By inducing *FAMA/FLP* under the *MUTE* promoter results in decreased amplitude of CDKs/CYCs pulse, leading to absence of final cell division. (D) Model diagram. (E and F) Experimental perturbation. Precocious expressionsd of *FAMA* (E) and *FLP* (F) during meristemoid-to-GMC transition. Shown are 7-day-old abaxial cotyledon epidermis expressing *MUTEpro::FAMA* (C) and *MUTEpro::FLP,* both conferring stoma with single GCs (yellow asterisks). Scale bars, 20 μm. Bottom insets, Mature GC GFP mature expression. (G) Quantitative analysis of SCGs in three independent transgenic lines expressing *FAMA* (n=14, 14, 14) and *FLP* (n=15, 15, 17) driven by the *MUTE* promoter. Values are mean ± s.e.m. (H) Mathematical modeling simulating the precocious expression of CDK/CYC in *fama* (reduced level of FAMA/FLP). When *FAMA/FLP* level is reduced and *CDK/CYC* is directly upregulated by *MUTE* promoter, the resulting *CDK/CYC* pulse amplitude is much higher than that of wild type. (I) Model diagram. (J) Perturbation experiments. Precocious expression of *CYCD5;1* in *fama* triggers supernumeral symmetric divisions. Shown are 2-week-old adaxial cotyledon epidermis of *mute* (top left), *fama* (top right), and *MUTEpro::CYCD5;1 fama* (bottom). The order of amplifying ACDs are numbered. (K) Quantitative analysis (violin plots) of cell numbers in each GMC tumors in *fama* (n= 43) and *MUTEpro::CYCD5;1 fama* (n=34). Dots, individual tumors; Pink rectangles, standard deviation. Wilcoxon rank sum test, p= 3.48e-08. (L) Persistent accumulation of *CYCD5;1* in the absence of *FAMA* or *FLP*. Confocal microscopy of GMC tumors in *fama* (top) and *flp-7* (bottom) expressing *CYCD5;1pro::CYCD5-GFP*.

Mathematical modeling of MUTE, FAMA/FLP and CDKs/CYCs faithfully reproduced the generation of single pulse of CDKs/CYCs observed *in vivo* (Figure 5B). We aimed for a minimal component modeling, which is qualitatively equivalent to three-component I1-FFL. All the interactions are implemented by Hill kinetics as described in original model with modifications (Mangan and Alon, 2003). Detail of the model is described in Supplemental materials & methods section.

To address the importance of the I1-FFL, we break up the I1-FFL *in silico* and experimentally verifying the stomatal phenotype as an outcome. If a node Y turns on simultaneously as X, the modeling predicts the output peak Z would decline to a sub-threshold level (Figure 5C). To test this prediction experimentally, *FAMA* as well as *FLP* were expressed precociously during the meristemoid-to-GMC transition driven by the *MUTE* promoter (Figure 5E-G). Both *MUTEpro::FLP* and *MUTEpro::FAMA* conferred differentiation of single-celled stomata (∼20% and ∼88%, respectively), expressing mature GC GFP marker (Figure 5E-G). The phenotype highly resembles that of the dominant-negative *CDKB1;1* and *CDKA1;1* as well as higher-order loss-of-function mutants of CDKB1s and CYCAs (Boudolf et al., 2004; Yang et al., 2014). In contrast, the previous report found no stomatal phenotype in *FLP* overexpressors (Lai et al., 2005). Our finding that *MUTEpro::FLP* triggers differentiation of single-celled stomata underscores the importance of specific cell type or developmental windows for *FLP* to function.

We next performed both simulation and experiments to break down the I1-FFL by turning on Z (*CDKs/CYCs*) simultaneously as X (*MUTE*), in which case the repression by Y (*FLP/FAMA*) would be too late to properly terminate the activity of Z (Figure 5H, I). We predicted that the repression by FLP/FAMA may be too strong to convincingly unravel the perturbed effects of *CYCD5;1*. We thus introduced *MUTEpro::CYCD5;1* into *fama* mutant background, which gives rise to GMC tumors with extra symmetric divisions (Figure 1J)(Ohashi-Ito and Bergmann, 2006). A precocious expression of *CYCD5;1* at the onset of *MUTE* expression triggered striking supernumeral divisions of *fama* GMC-like tumors, vastly increasing the number of cells per tumor (Figure 5J, K). The finding corroborates with the mathematical modeling (Figure 5H).

As shown in Figure 3A, CYCD5;1 disappears before the SCD of GMCs. To further address whether FAMA and FLP are required for the repression of CYCD5;1, we introduced CYCD5;1-GFP into *fama* and *flp-7* mutant backgrounds. Indeed, strong CYCD5;1-GFP signals are accumulated in symmetrically-dividing GMC tumors (Figure 5L). Combined, both mathematical modeling and experimental perturbations demonstrate the critical, direct role of MUTE in orchestrating the regulatory feed-forward loop ensuring the one symmetric division to create a stoma.

### Saturation and noise in ectopic *iMUTE* could flip the outcome of I1-FFL

Our mathematical and experimental analyses revealed that regulatory motif controlled by MUTE enables a fast response time and transient upregulation of cell-cycle regulator gene expression. Previous studies reported that *MUTE* overexpression confers constitutive stomatal differentiation in the cotyledon/leaf epidermis (MacAlister et al., 2007; Pillitteri et al., 2007; Trivino et al., 2013). However, through characterizing of our model parameters, we found that MUTE has to regulate *FAMA/FLP* much more tightly to ensure the single SCD under *iMUTE* overexpression (Figure 6A, B). Here, sustained *iMUTE in silico* limits the possible range of strong CDKs/CYCs activation by MUTE. On the other hand, *iMUTE* overexpression causes stronger activation of *FAMA/FLP*. In the parameter sets we employed, this could lead to faster decline of CDKs/CYCs, diminishing the peak below a threshold to trigger the SCD. Taking into account the modeling results that predict the dysregulation of the MUTE-orchestrated I1-FFL, we sought to revisit the *MUTE* overexpression phenotype.

**Figure 6.**
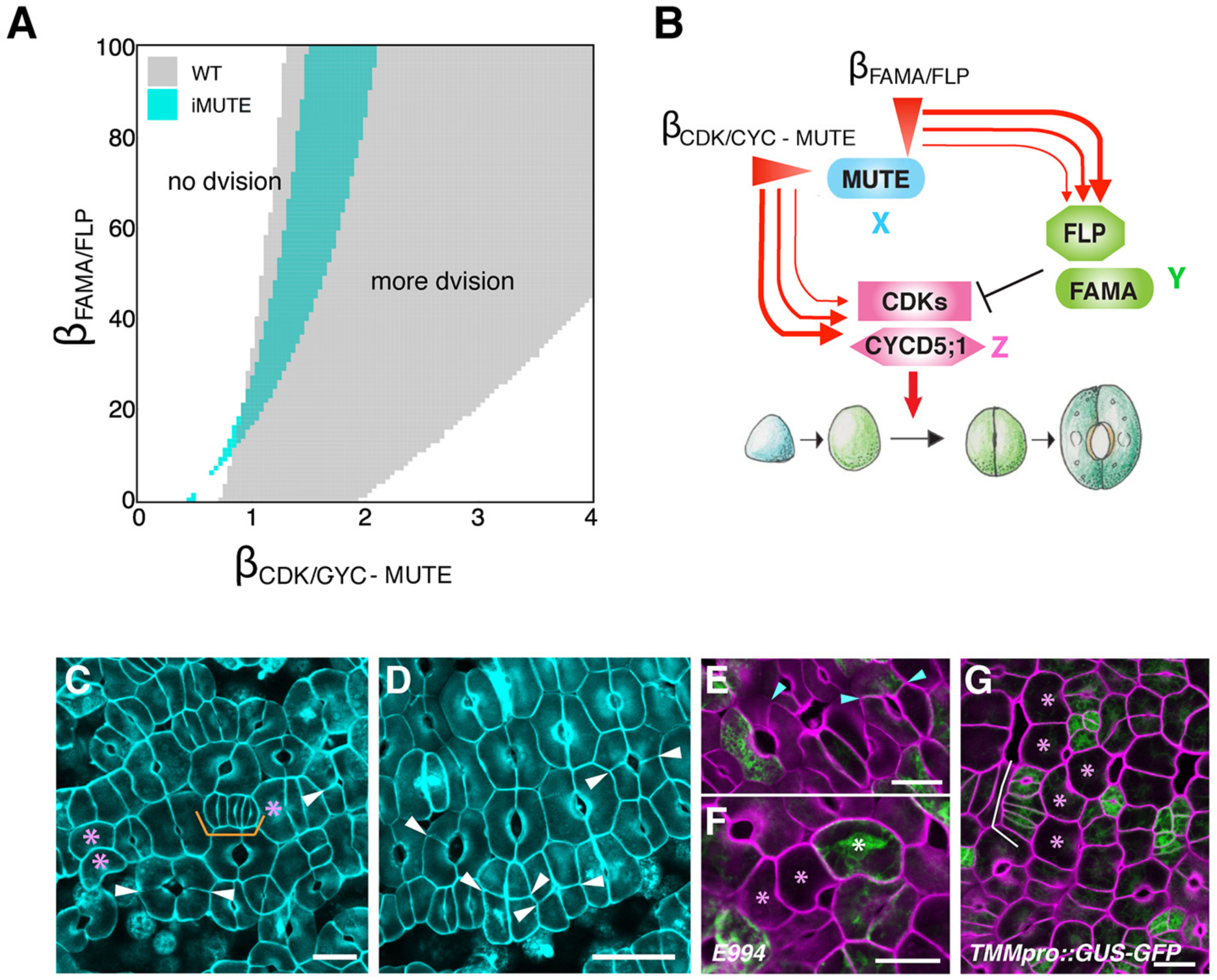
Sustained, saturated MUTE expression could result in variable outcomes of stomatal differentiation. (A) Numerical simulation results obtained by changing the strength of the effect of MUTE over FAMA/FLP and CDKs/CYCs expression. The results were classified based on the duration of CDKs/CYCs activation. The parametric regions corresponding to the single cell division in wild type (gray) and *iMUTE* (cyan). (B) Schematic Diagram showing the different strengths of MUTE on FAMA/FLP and CDKs/CYCs under *iMUTE* overexpression. (C-G) Cotyledon abaxial epidermis of 3-day-old *iMUTE* seedlings grown in the presence of estradiol. Each image was taken from individual seedling. While most epidermal cells differentiate into stomata, some become singular GCs (B, E, F, pink asterisks), rows of stomata from parallel extra divisions of GMCs (B, orange bracket; F, white bracket), or 3-4 celled stomata (white arrowheads) A subset of 3-4 celled stomata (D, plus) and singular GCs (E, white asterisk) express mature GC GFP marker, whereas parallel-dividing GMCs retain stomatal-lineage marker *TMMpro::GUS-GFP* (F, white bracket). Scale bars, 20 μm.

Indeed, careful observations of *iMUTE* epidermis revealed that, within the sheet of stomata-only epidermis, occasionally formed are singular GCs, *fama*-like GMC tumors, and stomata made with a trio or quartet of GCs surrounding a pore (Figure 6C-G). The singular GCs (Figure 6C,F,G, pink asterisks) are the hallmark of *FAMA* overexpression (Ohashi-Ito and Bergmann, 2006), whereas the excessive symmetric divisions (Figure 6C,G, orange and white brackets) imply the loss of *FAMA* or *FLP* activities. Conversely, 3-4 celled stomata (Figure 6C,D, white arrowheads) are the signature of ectopic activities of cell cycle genes in GMCs (Adrian et al., 2015; Yang et al., 2014). Mature GC GFP marker was expressed in a subset of GCs in 3-4 celled stomata (Figure 6E, cyan arrowheads) and likewise in a subset of singular GCs (Figure 6F, white asterisks). Thus, regardless of developmental outcome as singular GCs or 3-4 celled stomata, *iMUTE* can trigger eventual GC differentiation. The supernumerary GMCs expressed stomatal-lineage GFP marker, *TMMpro::GUS-GFP* (Figure 6G, white bracket), corroborating their identity.

## Discussion

This study identifies the complete inventories of early MUTE-responsive genes. The comparison of SPCH and MUTE shared and unique targets revealed how MUTE switches the cell-cell signaling from stomatal lineage initiation to commitment. The work further unraveled that MUTE directly induces the expression of both the cell cycle regulators and their transcriptional repressors, thereby orchestrating the I1-FFL to generate the robust single symmetric division event to create functional stomata.

### Logic of cell-state transition by sequential actions of bHLH proteins

How could later-acting stomatal bHLHs switch the precursor state from their earlier acting sisters? Our study revealed that MUTE binds to the SPCH-binding sites of the shared target genes and takes over their lineage-specific expressions, while repressing the earlier-acting gene *EPF2* to switch cell-cell signaling circuits (Figure 1L). The regulatory modules of stomatal differentiation resemble that of myogenesis in animals, where myogenic bHLHs; Myf5, MyoD, Myogen and MRF4 sequentially direct lineage specification, proliferation, and differentiation (Putarjunan and Torii, 2016; Tapscott, 2005). Extensive ChIP-seq studies of Myf5 and MyoD have shown that these two myogenic bHLHs bind to the nearly identical target sites genome-wide. However, unlike Myf5, MyoD promotes strong transcriptional activation via Pol II recruitment, suggesting that their functional specificities lie in their transcriptional activities (Conerly et al., 2016). Each stomatal bHLH possesses a unique motif signifying its function (Davies and Bergmann, 2014). Interestingly, overexpression of SPCH without the MAP kinase target domain or truncated FAMA lacking the N-terminal activation domain phenocopies MUTE activities (Lampard et al., 2008; Ohashi-Ito and Bergmann, 2006), suggesting that these additional modules prevent the functional interference among the three bHLHs.

It is known that *SPCH, MUTE*, and *FAMA* expressions are tightly regulated by the epigenetic mechanisms (Lee et al., 2014; Matos et al., 2014). The local chromatin state may explain why some targets (e.g. *TMM, SCRM*) are immediately induced by *MUTE* while others (e.g. *FAMA*) delay for ∼8 hrs. It could also explain the previous report that the ability for *MUTE* to induce stomatal differentiation becomes restricted as plants age (Trivino et al., 2013). In myogenesis, both Myf5 and MyoD recruit histone acetyltransferase to alter the epigenetic landscape at their target sites (Cao et al., 2010; Conerly et al., 2016). It would be interesting to test in future whether local and global epigenetic landscapes are regulated by each stomatal bHLH.

### MUTE as a potent inducer of cell division

Our study unraveled that MUTE is a potent inducer of cell cycle genes (Figure 2). MUTE strongly upregulates *CDKBs* (*CDKB1;1* and *CDKB1;2*) and *CYCA2s* (*CYCA2;2*, *CYCA2;3*) that promote GMC symmetric divisions (Boudolf et al., 2004; Xie et al., 2010). CDKBs-CYCA2s complexes are known to regulate S/G2 phase, but do not drive the cell cycle entry. Our work further identified CYCD5;1 as a D1-cyclin promoting the symmetric division. *CYCD5;1* is known to partner with CDKA1;1 (Boruc et al., 2010), which is not likely a MUTE target (Figures 2, S5). Because G1/S transition is a rate-limiting step, once *CYCD5;1* expression is induced, basal levels of CDKs and G2/M cyclins in *mute* may be sufficient to execute the symmetric-division-like cell division. Time-lapse imaging shows that CYCD5;1 peaks and disappears ∼8 hrs before the symmetric division prior to CDKB1;1 accumulation (Figure S6). The sequential peaks of CYCD5;1 followed by CDKB1;1 are consistent with their roles in G1/S and G2/M transitions, respectively.

It is worth noting that modest enrichments of *CDKB1;1* and *CYCA2;3* were reported in *scrm-D mute* mutant, which does not execute the symmetric division (Pillitteri et al., 2011). Because CDKB1;1 and CYCA2s suppress endocycles (Boudolf et al., 2009), it is possible that these cell cycle genes exhibit background-level expressions in the MUTE-independent manner, which may be crucial for preventing the endoreduplication of stomatal-lineage cells. In this scenario, the primary role of MUTE is to boost their timely expressions above the threshold level in order to drive the symmetric division. In this regard, it is interesting that CYCD5;1 has been reported as a candidate quantitative trait gene for endoreduplication in Arabidopsis natural accessions (Sterken et al., 2012). In any event, Arabidopsis MUTE as a potent inducer of cell division accords with the role for its mobile Brachypodium ortholog, BdMUTE, in promoting the subsidiary cell division (Raissig et al., 2017). Whether BdMUTE (or other grass MUTE orthologs) directly drives the symmetric division of grass stomata is a future question.

### I1-FFL orchestrated by MUTE drives the single symmetric division to create stomata

Our study unraveled that MUTE directly activates the expressions of cell-cycle genes and the direct repressors of the cell cycle genes. Furthermore, our modeling showed that the I1-FFL orchestrated by MUTE can trigger a single pulse of gene expression, in this case the cell cycle genes, within the narrow developmental windows encompassed by MUTE and FAMA/FLP (Figure 5). Importantly, the single pulse is much more robustly generated by the endogenous, pulsed *MUTE* expression than for saturated and sustained one (Figure 6). The I1-FFL is known to function as a pulse generator (Basu et al., 2004; Mangan and Alon, 2003): the circuit can generate a pulse output even under sustained input. This explains why sustained *iMUTE* overexpression still largely produces ‘normal’ stomata with paired GCs. Since *MUTE* expression window is limited in the wild type, theoretically, the simple linear circuit could be implemented for a pulse output. However, the I1-FFL would hold advantages for this biological context. The I1-FFL can accelerate the response time (Mangan and Alon, 2003), thus allowing MUTE to achieve the single division event concomitantly with stomatal differentiation. Our model is consistent with a previous report for a step input (Goentoro et al., 2009) such that a delay in the response of *FAMA/FLP* to MUTE enabled the amplitude and duration of *CDKs/CYCs* activation to be increased, which contributes to the sharp and high peak. This emphasizes the importance of the direct control of FAMA/FLP by MUTE to achieve such coordination. Recently, the I1-FFL was implicated in transcriptional control of root Casparian strip differentiation (Fernandez-Marcos et al., 2017). Thus, plants may use I1-FFL for critical cell-fate decision-making processes in broader contexts.

## Acknowledgements

We thank L. Pillitteri for generating *iMUTE* lines; E. Lee, F. Sack, S. Vanneste, D. Inzé, L. de Vylder, and J. Garrick for mutants and reporter lines; C. Zheng for plant care; M. Ito for expert comments on the manuscript. The work was supported by the US National Science Foundation (MCB-0855659) and the Gordon and Betty Moore Foundation (GBMF-3035) to K.U.T, who acknowledges support from HHMI and University of Washington.

## Author contributions

Conceived the project, K.U.T.; designed experiments, S.K.H. and K.U.T.; performed experiments, S.K.H., X.Q., J.H.D., K.L.M., E.K. K.U.T.; performed bioinformatics analysis, S.K.H. and T.A.E.; analyzed data, S.K.H, X.Q., E.K., T.A.E, and K.U.T.; performed mathematical modeling; K.S., T.M.; Wrote the paper, K.U.T; All authors contributed to finalizing the paper.

