## Supplementary Materials for "MUTE Directly Orchestrates Cell State Switch and the Single Symmetric Division to Create Stomata"

### EXPERIMENTAL MOEDL AND SUBJECT DETAILS

The Arabidopsis Columbia (Col) accession was used wild type. The following mutants and reporter transgenic lines were reported previously: *mute*, *fama*, *MUTEpro::MUTE-GFP, FAMApro::FAMA-GFP* (Pillitteri et al., 2007); *mute-2* (Pillitteri et al., 2008); *scrm-D* (Kanaoka et al., 2008); *TMMpro::GUS-GFP* (Nadeau and Sack, 2002); *E994* (Pillitteri et al., 2007); *PM-RB* (Pillitteri et al., 2011); *iMUTE* (Qi et al., 2017); *flp-7* and *FLPpro::GUS-GFP* (Lai et al., 2005). The original *flp-7* allele (Lai et al., 2005) is in Landsberg *erecta* (L*er*) background. We have outcrossed *flp-7* to Col accession for three times to remove the *er* mutation, which affect stomatal development, and clean up the genotype. Reporter lines were introduced into respective mutant backgrounds by genetic crosses or Agrobacterium-mediated transformation (see below for methods and Key Reources Table for the lines generated) and genotypes confirmed by PCR. For sequence of genotyping primers, see Table S4. Seedlings and plants were grown as described previously (Lee et al., 2012).

### QUANTIFICATION AND STATISTICAL ANALYSIS

The Leica LAS AF software and Imaris ver. 8.1.3 (Bitplane) were used for post-acquisition image processing. Quantitative analysis of MUTE signal intensity was performed using Imaris ver. 8.1.3 briefly as described previously (Qi et al., 2017). Briefly, an extensive series of Z-stack confocal images covering the entire meristemoids (~17 layers) were subjected to surface rendering in green channel to capture the *CYCD5;1pro::CYCD5;1-GFP* and *CYCD5;1pro_Δ3E-box::CYCD5;1-GFP* signals in the 3-D space. Cut-off value was set 0.85 for sphericity. Intensity sum and standard deviation for each nucleus were subsequently calculated. Statistical analyses were performed using R ver. 3.3.1. The Wilcoxon Rank Sam Test was performed when comparing guard cell numbers (e.g. *fama* tumors) in individual stomata/stomatal precursors as well as for GFP quantification, because normal distribution among the population may not be expected. Graphs were generated using R ggplot2 package or Microsoft Excel.

### DATA AND SOFTWARE AVAILABILITY

The RNA-seq data generated in this study are deposited to the NCBI Gene Expression Omnibus (GEO) with an accession number: GSE107018. All codes for our mathematical modeling, written in Mathematica (Wolfram, Inc), is available as a Supplementary Document (Document S1). For software for bioinformatics, image-analysis, and mathematical modeling, see Key Resource Tables.

### METHOD DETAILS

#### Plasmid Construction and Transgenic Plants Generation

The following constructs were generated for the study: pLJP245 (*Est::MUTE [iMUTE]*), pLJP246 (*Est::SPCH [iSPCH]),* pSKH198 *(CYCD5;1pro::gCYCD5;1-GFP),* pSKH238 *(CYCD5:1pro_Δ3E::gCYCD5;1-GFP),* pSKH128 *(MUTEpro::gCYCD5;1),* pSKH133 *(MUTEpro::gFLP),* pSKH137 *(MUTEpro::gFAMA)* and pJT164 *(FAMApro::FAMA-RFP)* For detailed information of each plasmid, see Supplementary Table S4. For generating transgenic Arabidopsis, plasmid constructs were electroporated into Agrobacterium (GV3101/pMP90) and subsequently transformed by floral dipping. At least three lines were subjected to detailed characterizations.

#### Estradiol-induction, RNA preparation, and RNA sequencing

Homozygous transgenic lines for *iMUTE* or *iSPCH* were grown on ½ MS agar media for 4 days. 40-50 seedlings for each replicate were bathed in a ½ MS liquid media containing 10 $\mu$M estradiol (Sigma-Aldric, E2758) or DMSO (mock). For the RNA-seq samples, three replicates per treatment (mock or estradiol) were collected 8hours after treatment for total RNA isolation. Three different batches of seedlings were harvested at the same time of the day to minimize the effects of circadian clock. Total RNA was isolated from the seedlings using RNeasy Plant Mini Kit (Qiagen, 74904) and DNase I digestion (Qiagen, 79254) was performed on column during the RNA extraction according to the instructions of the manufacturer. The RNA samples were sent to the New York Genome Center (NYGC) facility for library construction and sequencing. RNA samples were run on the Agilent 2100 bioanalyzer to access the RNA quality. mRNA libraries were prepared by using Illumina TruSeq Stranded mRNA and read paired-end 50bp in length with 30M coverage per sample on Illumina Hiseq 2500 system.

**Bioinformatics and Data Analysis**

Sequences of RNA-seq experiments, those performed in this study and those obtained from a public database (GSE57953, GSE57497 and GSE58856, NCBI GEO), were mapped on TAIR 10 genome using TopHat2-PE (ver. 2.0.13, https://github.com/infphilo/tophat) with default options, and the Illumina reads were mapped and the read counts were calculated using our in-house programs with the gene location GFF file distributed at TAIR (ftp://ftp.arabidopsis.org/Genes/TAIR10_genome_release/TAIR10_gff3/TAIR10_GFF3_genes.gff). Differentially expressed genes and their statistical significance were evaluated using DESeq2 R package (http://bioconductor.org/packages/release/bioc/html/DESeq2.html). Upregulated and downregulated genes in RNA-seq experiments were defined as log2 (treated/control) more than 0.5 or less than 0.5, and q-values less than 0.05. Qualities and reproducibility of our *iMUTE* induced and mock control triplicate samples were verified by evaluating correlations between logarithms of RPKM (read per kilobase exon per million mapped reads). Values of Pearson's correlation were more than 0.980 between samples of the same condition (control vs control or treated vs treated of the three biological replicates) and more than 0.946 between different conditions. The published FACS-sorted stomatal cell-lineage transcriptome samples (Adrian et al., 2015) showed low reproducibility, and none of the biolgical replicates of stomatal-lineage specific populations clustered correctly (Fig. S1). Likewise, the published inducible SPCH (iSPCH) and control wild-type RNA-seq data (Lau et al., 2014) showed low reproducibility (Fig. S1). While our data are triplicated pair-end experiments and the others were duplicated/triplicated single-end experiments (Lau et al., 2014; Adrian et al., 2015), we assumed that the difference did not affect much for the scope of this study.

A ChIP-seq experiment using anti GFP antibody (GSE57953) was also downloaded from NCBI GEO and aligned on TAIR 10 genome using bowtie2 program (version 2.1.0, http://bowtie-bio.sourceforge.net/bowtie2/) with default parameters. Genes were classified for some functional groups using the annotation obtained from the Gene Ontology database (http://geneontology.org, validation date 9/1/2016) and literatures. Stomatal genes were defined as genes having “stomatal complex morphogenesis (GO:0010103)” and/or “stomatal complex development (GO:0010374)” and curated using the literature (Horst et al., 2015; Lau et al., 2014; Peterson et al., 2013; Pillitteri et al., 2011). Cell cycle, cell division, mitotic and auxin-related genes were defined as all genes having GO annotations that contained “cell cycle,” “cell division," “mitotic” and “auxin” such as GO:0007049 (cell cycle), GO:0051726 (regulation of cell cycle), GO:0051301 (cell division), GO:0051302 (regulation of cell division), GO:0007067 (mitotic nuclear division), GO:0006312 (mitotic recombination), GO:0009733 (response to auxin), and GO:0009734 (auxin-activated signaling pathway).

To cluster the expression patterns, we calculated logarithm of odds score (LOD) of contingency tables for upregulated genes between two experiments in order to minimize biologically-irrelevant clustering based on the type of experimental procedures (e.g. minimize clustering on the basis of an upregulation of stress signatures due to protoplasting followed by a FACS sorting [Adrian et al., 2015]). The experiments were clustered using a hierarchal approach, in which the pairs having most score were clustered into a new virtual node until no pairs exist. Heat maps and Venn diagrams were generated using in house python scripts.

**cDNA preparation and qRT-PCR**

For time-course induction, estradiol-inducible MUTE seeds were sown on 1/2 media, and subjected to stratification at 4°C for two days then grown at the regular long-day condition for three or four days. Subsequently, 40-50 seedlings were transplanted to 1/2 MS media containing DMSO (mock) or 10 µM estradiol (*iMUTE*) and sequentially harvested after 2, 4, 6, 8, 10, 12 hours after induction. To reduce the effects of circadian clock, seedlings were transferred at the same time of the day. 1 µg or RNA was converted to cDNA using iScript cDNA synthesis kit (Bio-Rad) according to the instructions of the manufacturer. First-strand cDNA was diluted to seven fold in double distilled water and used as template for Real-time qPCR. qRT-PCR was performed as described in (Qi et al., 2017). At least four 10-fold dilution series were amplified by the same primers to set standard curve and calculate the quantity of the gene expression. For each experiment, three technical replicates were performed, and for all experiments at least 3 biological replicates were performed. Relative expression was calculated by dividing *ACTIN2* gene expression over the specific-gene expression. See Table S4 for primer sequences used for qRT-PCR.

### Chromatin Immunoprecipitation

Total 160 mg (2 plates) of seeds were sown on ½ MS agar media, and seedlings at 42~48 hrs after germination were collected and snap frozen in liquid nitrogen. Frozen tissue was ground to fine powder and re-suspended in 20ml of ChIP extraction buffer 1 (Bowler et al., 2004) containing 1% formaldehyde (Sigma, F1635), and was incubated at 4 °C for 10 min for DNA-protein cross-linking. 2 M glycine was added to quench the reaction, then samples were incubated at 4 °C for 5 min. Subsequent shearing of DNA using Bioruptor Plus UCD-300 sonicator (Diagenode), immunoprecipitation using Dynabeads protein G (Invitrogen), as well as capturing, washing, reverse cross-linking of immune-complex and DNA purification- were performed as described previously (Qi et al., 2017). Quantitative PCR (qPCR) was carried out using gene specific primers (Table S4). The qPCR was run using iTaq Universal SYBR Green Supermix on CFX96 real-time system (Bio-Rad) as decribed previously (Qi et al., 2017). At least three technical replicates were performed for each experiment. For each experiment, at least three biological replicates were performed. Dilution series of input was amplified simultaneously by the same primers to calculate the quantity of the ChIPed DNA as % input (Yamaguchi et al., 2014).

### Microscopy

For confocal microscopy, cell peripheries of seedlings were visualized with either propidium iodide (Molecular Probes). Images were taken using LSM700 (Zeiss) for GFP as described previously or SP5 (Leica) was used with White Light Laser (excitation at 488 nm and emission at the 500-515 nm for GFP; excitation 619 nm and emission at 642 nm for propidium iodide) using HyD detector. The confocal images were false colored, and brightness/contrast were uniformly adjusted using Photoshop CS6 (Adobe).

For time-lapse live imaging, Arabidopsis germinating cotyledons expressing dual reporters (plasma-membrane RFP with either *CYCD5;1pro::CYCD5;1-GFP*, *MUTEpro::MUTE-GFP*, or *FAMApro::FAMA-GFP*) were generated by genetic crosses and subjected to time-lapse imaging with 30-min intervals as described previously (Peterson and Torii, 2012; Qi et al., 2017).

### Mathematical modeling

We formulated the interaction of MUTE, FAMA/FLP, and CDKs/CYCs as type 1 incoherent feed-forward loop model (Mangan and Alon, 2003). In this model, time course of MUTE expression level ([*MUTE*]) is given, and the resulting FAMA/FLP and CDKs/CYCs expression ([*FAMA/FLP*] and [*CDKs/CYCs*], respectively) are under the control of MUTE expression. For minimal component, we hereby combine CDKs/CYCs as one parameter. Thus, the model does not account for specific cell cycle phase. Likewise, both FAMA and FLP are simplified as one parameter. We also introduced MUTE promoter activity as [*MUTEpro*].

We define wildtype MUTE promoter activity as

$$[MUTEpro](t) = \begin{cases} 1 & (0 < t < 5) \\ 0 & (\text{otherwise}) \end{cases}.$$

In the iMUTE mutant, the promoter activity pattern is defined as follows:

$$[MUTEpro](t) = \begin{cases} 2 & (0 < t) \\ 0 & (\text{otherwise}) \end{cases}.$$

We assume MUTE expression is controlled by MUTE promoter activity as follows:

$$\frac{d[MUTE]}{dt}(t) = \beta_{MUTE}[MUTEpro](t) - \alpha_{MUTE}[MUTE](t).$$

$\beta_{MUTE}$ represents the strength of the effect of MUTE promoter over MUTE expression. $\alpha_{MUTE}$ is the decay rate of MUTE. FAMA/FLP expression is assumed to be controlled by MUTE as follows:

$$\frac{d[FAMA/FLP]}{dt}(t) = \beta_{FAMA/FLP}\, f_a([MUTE](t-\tau), K) - \alpha_{FAMA/FLP}[FAMA/FLP](t).$$

$\beta_{FAMA/FLP}$ represents the strength of the effect of MUTE over FAMA/FLP expression. $\alpha_{FAMA/FLP}$ is the decay rate of FAMA/FLP. $f(u, K)$ represents the effect of a gene on another gene expression. Activation reaction is defined by

$$f_a(u, K) = \frac{u}{K + u},$$

and repression reaction is defined by

$$f_r(u, K) = \frac{K}{K + u}.$$

$\tau$ is the delay of the effect of MUTE over FAMA/FLP. This delay is introduced based on the induction kinetics of *FAMA* transcripts upon estradiol induction of *iMUTE*.

CDKs/CYCs expression is controlled both by MUTE and FAMA/FLP as follows:

$$\begin{aligned}\frac{d[CDKs/CYCs]}{dt}(t) &= B_{CDKs/CYCs} + \beta_{CDKs/CYCs-MUTE}\, f_a([MUTE](t), K_{MUTE}) \\ &+ \beta_{CDKs/CYCs-FAMA/FLP}\, f_r\big([FAMA/FLP](t), K_{FAMA/FLP}\big) \\ &- \alpha_{CDKs/CYCs}[CDKs/CYCs](t).\end{aligned}$$

$B_{CDKs/CYCs}$ represents the basal production or reduction of CDKs/CYCs. $\beta_{CDKs/CYCs-MUTE}$ and $\beta_{CDKs/CYCs-FAMA/FLP}$ are the strength of the effect of MUTE and FAMA/FLP over CDKs/CYCs expression. $\alpha_{CDKs/CYCs}$ is the decay rate of CDKs/CYCs. We incorporate the effects of MUTE and FAMA/FLP over CDKs/CYCs additively, not synergistically. If there is the basal reduction of

CDKs/CYCs, i.e. $B_{CDKs/CYCs} < 0$ , we also assumed $[CDKs/CYCs](t)$ as 0 if it became less than 0.

For MUTEpro::FLP and MUTEpro::FAMA, we incorporated direct induction of FAMA/FLP by MUTE promoter as follows:

$$\frac{d[FAMA/FLP]}{dt}(t) = \beta_{FAMA/FLP}\, f_a([MUTE](t-\tau), K) + \beta_{MUTEpro-FAMA/FLP}[MUTEpro](t) - \alpha_{FAMA/FLP}[FAMA/FLP](t).$$

$\beta_{MUTEpro-FAMA/FLP}$ represents the strength of the direct induction of FAMA/FLP by MUTE promoter. For *MUTEpro::CYCD5;1*, we incorporated direct induction of CDKs/CYCs by MUTE promoter as follows:

$$\frac{d[CDKs/CYCs]}{dt}(t) = B_{CDKs/CYCs} + \beta_{CDKs/CYCs-MUTE}\, f_a([MUTE](t), K_{MUTE}) + \beta_{CDKs/CYCs-FAMA/FLP}\, f_r\big([FAMA/FLP](t), K_{FAMA/FLP}\big) + \beta_{MUTEpro-CDKs/CYCs}[MUTEpro](t) - \alpha_{CDKs/CYCs}[CDKs/CYCs](t).$$

We reduced $\beta_{FAMA/FLP}$ for *fama* mutant background.

For the analysis of parametric dependence, we calculated $d$, the duration of the first continuous period satisfying $[CDKs/CYCs](t) > 3$. We assumed the single cell division is achieved when *d* satisfied $8 < d < 16$.

Numerical simulation was done with explicit Euler scheme with *Mathematica* (Wolfram, Inc., Champaign, IN). Complete source code for the numerical simulation is provided by supplementary data.

### REFERENCES TO EXPERIMENTAL PROCEDURES


Kanaoka, M.M., Pillitteri, L.J., Fujii, H., Yoshida, Y., Bogenschutz, N.L., Takabayashi, J., Zhu, J.K., and Torii, K.U. (2008). SCREAM/ICE1 and SCREAM2 specify three cell-state transitional steps leading to arabidopsis stomatal differentiation. Plant Cell *20*, 1775-1785.

Lai, L.B., Nadeau, J.A., Lucas, J., Lee, E.K., Nakagawa, T., Zhao, L., Geisler, M., and Sack, F.D. (2005). The Arabidopsis R2R3 MYB proteins FOUR LIPS and MYB88 restrict divisions late in the stomatal cell lineage. Plant Cell *17*, 2754-2767.

Lee, J.S., Kuroha, T., Hnilova, M., Khatayevich, D., Kanaoka, M.M., McAbee, J.M., Sarikaya, M., Tamerler, C., and Torii, K.U. (2012). Direct interaction of ligand-receptor pairs specifying stomatal patterning. Genes Dev *26*, 126-136.

Nadeau, J.A., and Sack, F.D. (2002). Control of stomatal distribution on the Arabidopsis leaf surface. Science *296*, 1697-1700.

Peterson, K.M., and Torii, K.U. (2012). Long-term, high-resolution confocal time lapse imaging of Arabidopsis cotyledon epidermis during germination. J Vis Exp.

Pillitteri, L.J., Bogenschutz, N.L., and Torii, K.U. (2008). The bHLH protein, MUTE, controls differentiation of stomata and the hydathode pore in Arabidopsis. Plant Cell Physiol *49*, 934-943.

Pillitteri, L.J., Sloan, D.B., Bogenschutz, N.L., and Torii, K.U. (2007). Termination of asymmetric cell division and differentiation of stomata. Nature *445*, 501-505.

Qi, X., Han, S.K., Dang, J.H., Garrick, J.M., Ito, M., Hofstetter, A.K., and Torii, K.U. (2017). Autocrine regulation of stomatal differentiation potential by EPF1 and ERECTA-LIKE1 ligand-receptor signaling. Elife *6*. e24102


### SUPPORTING INFORMATION

#### Figures S1-S6

**Figure S1. Reproducibility test of RNA-seq samples and qRT-PCR samples analyzed in this study (related to Figure 1).**

**Figure S2. GO categories highly enriched in *iMUTE* up or down-regulated genes and comparison with *iSPCH* (related to Figure 1).**

**Figure S3. Additional three biological replicates of qRT-PCR of stomatal genes (related to Figure 1).**

**Figure S4. Additional biological replicates of ChIP-qPCR assay and qRT-PCR (related to Figures 1, 2, 4).**

**Figure S5. Additional biological replicates of qRT-PCR of cell-cycle related genes (related to Figure 2).**

**Figure S6. Time-lapse imaging of CDKB1;1 and FAMA reporter protein accumulation during stomatal differentiation (related to Figure 3).**

**Tables S1-S5**

**Table S1. RNA-seq analysis of iMUTE, and re-analysis of iSPCH and stomatal lineage transcriptome (related to Figure 1).**

**Table S2. List of GO terms enriched in differentially expressed genes by iMUTE and iSPCH (related to Figures 1 and S2).**

**Table S3. Stomatal genes, Cell cycle, cell division, and mitotic (CC+CD+Mitotic) genes up-regulated by iMUTE (related to Figures 1 and 2).**

**Table S4. List of plasmids and primers.**

**Table S5. Parameters of mathematical modeling (related to Figures 5, 6)**

**Movies S1-S3**

**Movie S1. Time-lapse movie of CYCD5pro::CYCD5-GFP during stomatal differentiation (related to Figure 3)**

**Movie S2. Time-lapse movie of MUTEpro::MUTE-GFP during stomatal differentiation (related to Figure 3)**

**Movie S3 Time-lapse movie of FAMApro::FAMA-GFP during stomatal differentiation (related to Figure 3)**

**Document S1**

**Mathematica codes for modeling I1-FFL (related to Figures 5, 6)**

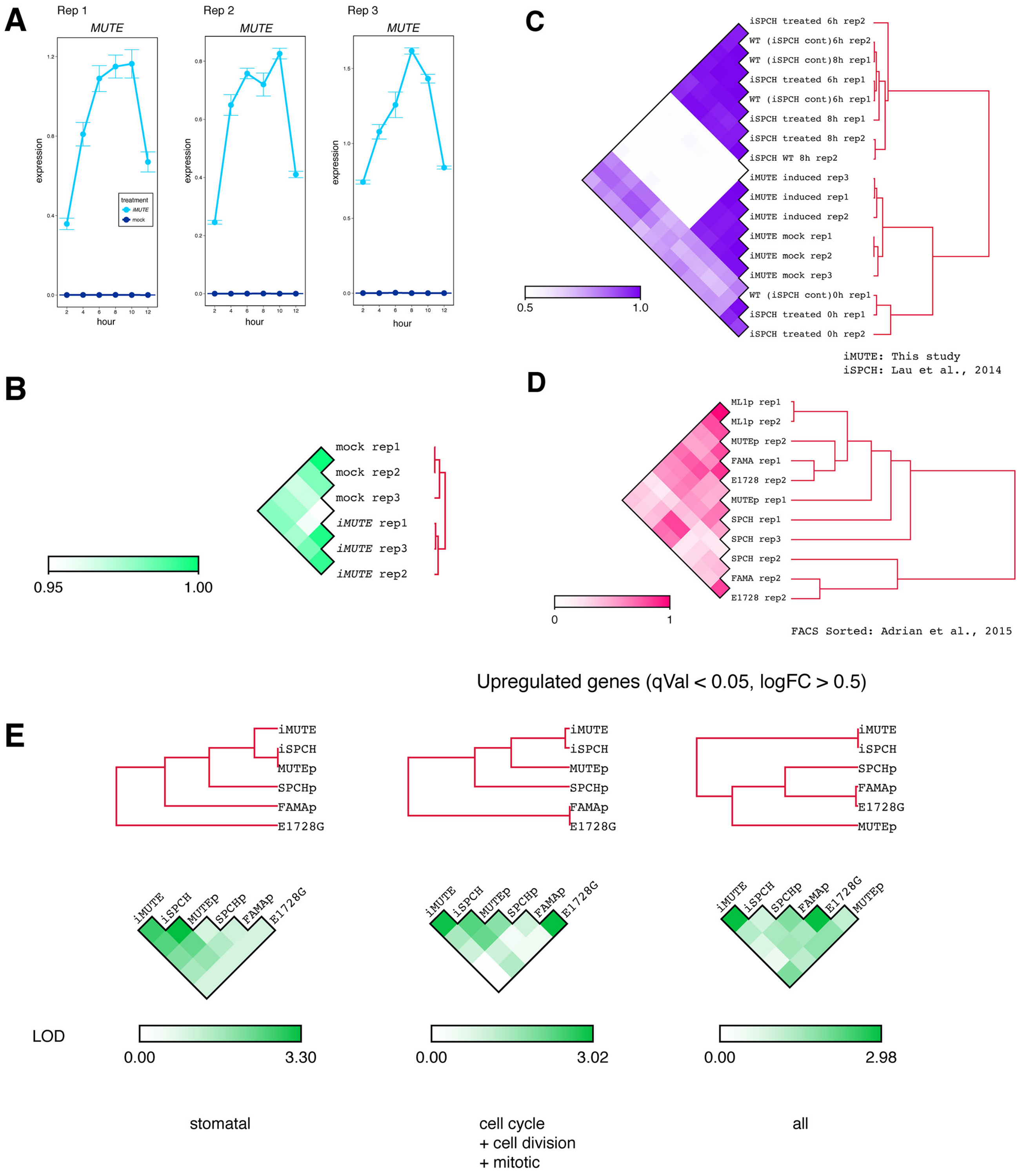
A
Rep 1
MUTE
Rep 2
MUTE
Rep 3
MUTE
expression
hour
treatment
iMUTE
mock
B
mock rep1
mock rep2
mock rep3
iMUTE rep1
iMUTE rep3
iMUTE rep2
0.95
1.00
C
iSPCH treated 6h rep2
WT (iSPCH cont)6h rep2
WT (iSPCH cont)8h rep1
iSPCH treated 6h rep1
WT (iSPCH cont)6h rep1
iSPCH treated 8h rep1
iSPCH treated 8h rep2
iSPCH WT 8h rep2
iMUTE induced rep3
iMUTE induced rep1
iMUTE induced rep2
iMUTE mock rep1
iMUTE mock rep2
iMUTE mock rep3
WT (iSPCH cont)0h rep1
iSPCH treated 0h rep1
iSPCH treated 0h rep2
0.5
1.0
iMUTE: This study
iSPCH: Lau et al., 2014
D
ML1p rep1
ML1p rep2
MUTEp rep2
FAMA rep1
E1728 rep2
MUTEp rep1
SPCH rep1
SPCH rep3
SPCH rep2
FAMA rep2
E1728 rep2
0
1
FACS Sorted: Adrian et al., 2015
Upregulated genes (qVal < 0.05, logFC > 0.5)
E
iMUTE
iSPCH
MUTEp
SPCHp
FAMAp
E1728G
iMUTE
iSPCH
MUTEp
SPCHp
FAMAp
E1728G
iMUTE
iSPCH
SPCHp
FAMAp
E1728G
MUTEp
LOD
0.00
3.30
0.00
3.02
0.00
2.98
stomatal
cell cycle
+ cell division
+ mitotic
all

**Figure S1. Reproducibility test of RNA-seq samples and qRT-PCR samples analyzed in this study.**

(A) Three biological replicates of qRT-PCR analysis of *MUTE* transcripts upon *iMUTE* induction. Transcript amounts were normalized against *ACTIN2* (*ACT2*).

(B) Pearson's correlation coefficients between three biological replicates of mock treated- and estradiol-induced *iMUTE* samples subjected to RNA-seq analysis (this study). High reproducibility (>0.98) among biological replicates.

(C) Person's correlation coefficiency analysis of the three biological replicates of iMUTE RNA-seq samples analyzed with the previously published *iSPCH* RNA-seq samples (Lau et al., 2014). The published wild-type controls (WT (iSPCH cont)) and *iSPCH* induced samples do not cluster together, implying low reproducibility of the biological replicates.

(D) Person's correlation coefficiency analysis of two biological replicates of the previously published FACS-sorted stomatal cell-state enriched transcriptomes (Adrian et al., 2015). The *AtML1* promoter (protodermal cell) RNAs cluster well, but other replicates do not cluster together, implying low reproducibility of the biological replicates.

(E) Logarithm of odds (LOD) score of contingency tables for upregulated genes between two experiments.

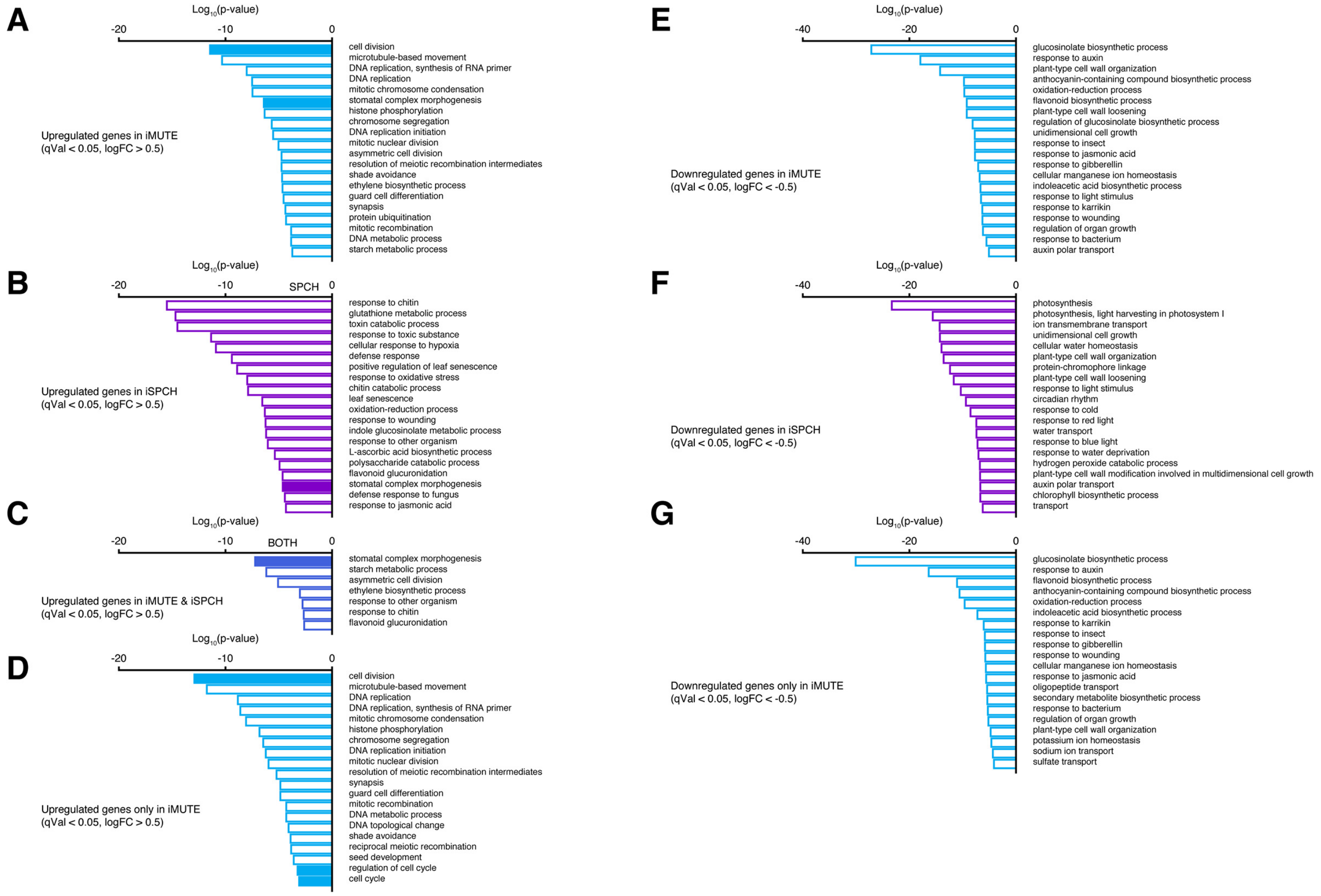


**Figure S2. GO categories highly enriched in *iMUTE* up or down-regulated genes and comparison with *iSPCH*.**

(A-D) GO categories and their p-values highly enriched in *iMUTE* (A), *iSPCH* (B) and both *iMUTE* and *iSPCH* (C) and in *iMUTE* only (D) up-regulated genes.

(E-G) GO categories and their p-values highly enriched in *iMUTE* (A), *iSPCH* (B) down-regulated genes and in *iMUTE* only down-regulated genes.

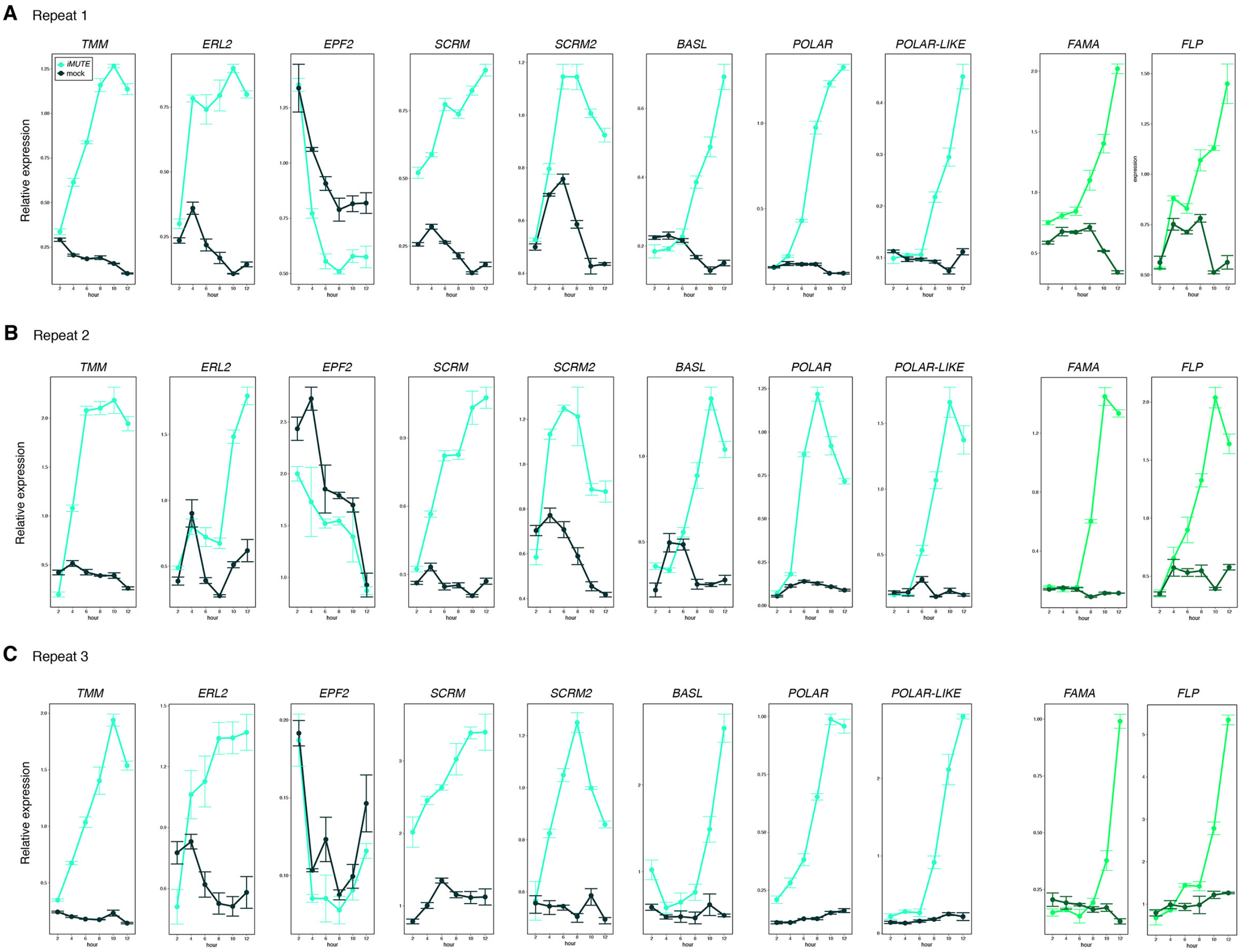


**Figure S3. Additional three biological replicates of qRT-PCR of stomatal genes.**

Quantitative RT-PCR analysis of the transcripts from 4-day-old seedlings that were either mock- or estradiol-treated for induced MUTE overexpression (*iMUTE*) at the time indicated. Relative expression represents qRT-PCR values normalized over that of *ACTIN2*. Values are Mean ± standard deviation of three technical replicates.

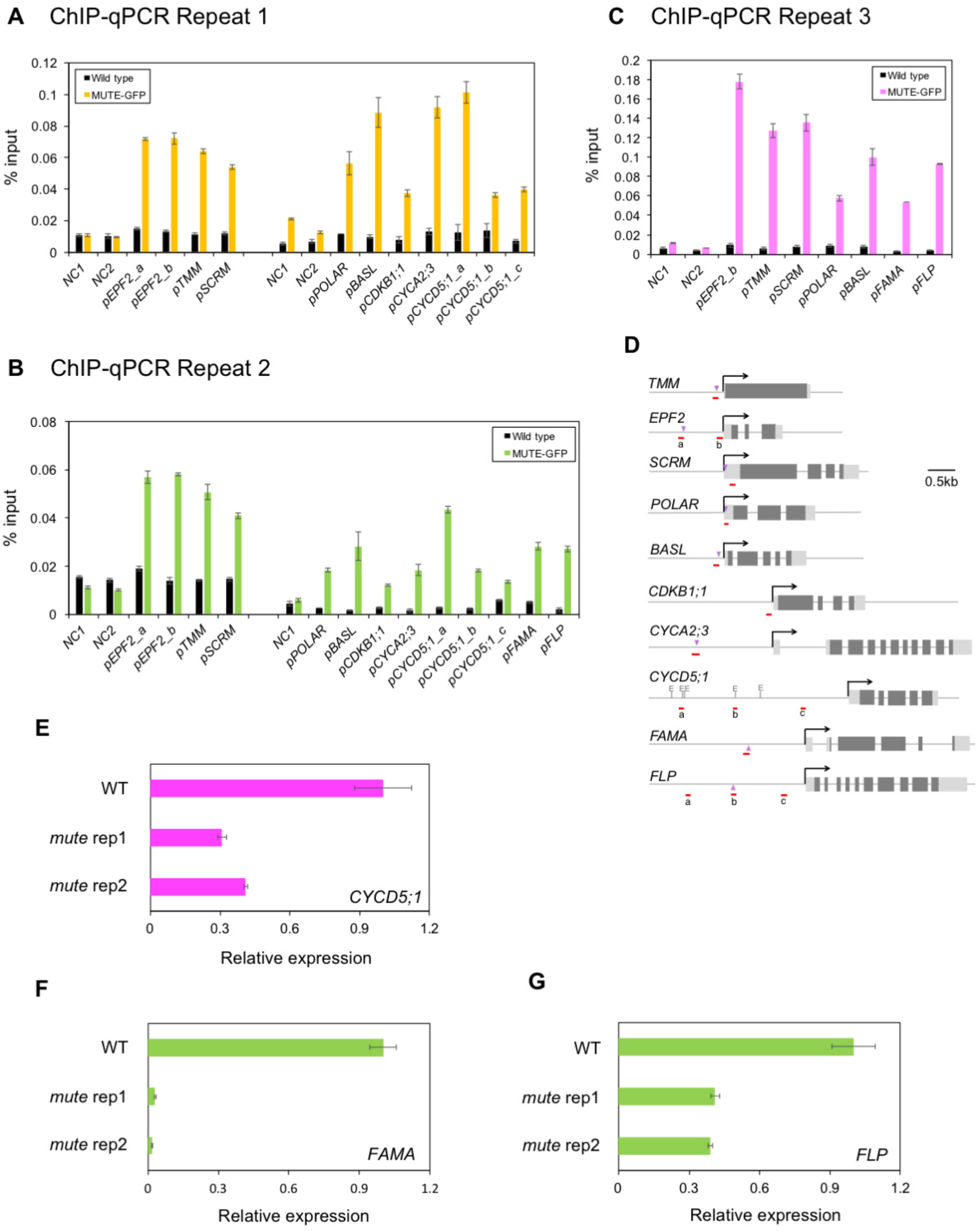

A ChIP-qPCR Repeat 1
% input
Wild type
MUTE-GFP
NC1
NC2
pEPF2_a
pEPF2_b
pTMM
pSCRM
pPOLAR
pBASL
pCDKB1;1
pCYCA2;3
pCYCD5;1_a
pCYCD5;1_b
pCYCD5;1_c
B ChIP-qPCR Repeat 2
pFAMA
pFLP
C ChIP-qPCR Repeat 3
D
TMM
EPF2
SCRM
POLAR
BASL
CDKB1;1
CYCA2;3
CYCD5;1
FAMA
FLP
0.5kb
E
WT
mute rep1
mute rep2
CYCD5;1
Relative expression
F
FAMA
G
FLP

**Figure S4. Additional biological replicates of ChIP-qPCR assay and qRT-PCR.**

(A-C) Quantitative PCR on the regions indicated in Fig. 1J, Fig. 2C and Fig. 4C after chromatin Immunoprecipitation (ChIP) against anti-GFP antibody in Col and transgenic plants expressing MUTE-GFP in *scrm-D*. Values are Mean ± S.E.M. of percent-input DNA of three technical replicates. (D) Diagrams of genes tested in ChIP-qPCR assay. Light gray rectangles, UTRs; dark gray rectangles, exons; arrows, transcriptional start sites; red line, amplicons; purple triangle, known SPCH binding sites (Lau et al., 2014); E, E-boxes. (E-G) Additional two biological replicates of quantitative RT-PCR (qRT-PCR) of *CYCD5;1* (E), *FAMA* (F) and *FLP* (G) transcripts in eight-day-old wild type (WT) and *mute* seedlings. The transcripts were normalized against *ACTIN2* and shown as expression fold change with transcripts in wild type normalized as 1.

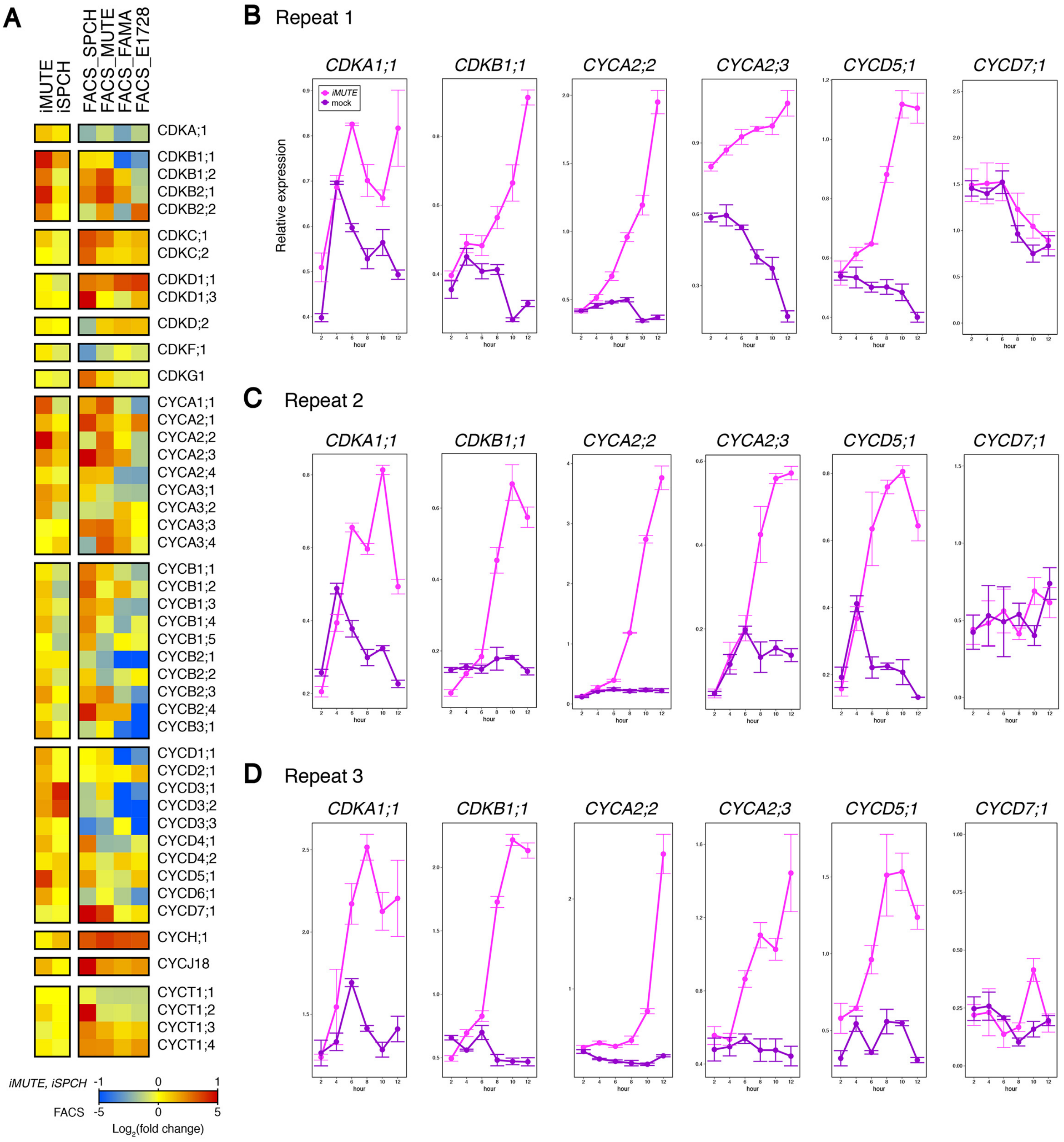


**Figure S5. Additional biological replicates of qRT-PCR of cell-cycle related genes.**

(A) Heat map showing fold change of expression of cell cycle genes by *iMUTE*, *iSPCH*, and FACS-sorted stomatal-lineage cells. (B-D) The conditions and methods for qRT-PCR is the same as described in Fig. S3 except for the seedling stage used in each replicate, 4-day-old seedling (B, D) and 3-day-old seedling (C) respectively. (E, F) Biological replicates qRT-PCR. Relative *CYCD5;1* expression in 7-day and 8-day-old wild-type and *mute* mutant seedlings (E) and relative *FAMA* and *FLP* expression in 8-day-old wild-type and *mute* seedlings. Bars, mean of three technical replicates. Error bars, mean ± s.d.

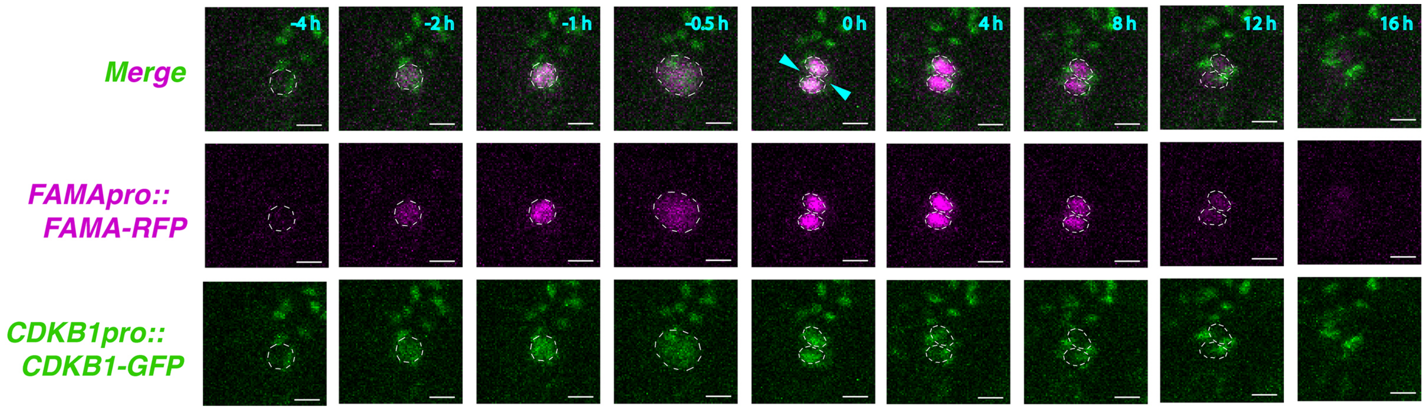


**Figure S6. Time-lapse imaging of CDKB1;1 and FAMA reporter protein accumulation during stomatal differentiation.**

Transient accumulation of CDKB1;1 (*CDKB1;1pro::CDKB1;1-GFP)* and FAMA (FAMApro::FAMA-RFP) during meristemoid-GMC-GC transition revealed by the time-lapse imaging. Cyan arrowhead indicates the single symmetric cell division (SCD) that gives rise to paired guard cells (hour 0 in cyan). Dotted circle indicates an outline of nucleus. Scale bar, 10 μm.
